# MRCK-1 activates non-muscle myosin for outgrowth of a unicellular tube in *Caenorhabditis elegans*

**DOI:** 10.1101/2024.01.25.577256

**Authors:** Evelyn M. Popiel, W. Brent Derry

## Abstract

The formation and patterning of unicellular biological tubes is essential for metazoan development. It is well established that vascular tubes and neurons use similar guidance cues to direct their development, but the downstream mechanisms that promote the outgrowth of biological tubes are not well characterized. We show that the conserved kinase MRCK-1 and its substrate the regulatory light chain of non-muscle myosin, MLC-4, are required for outgrowth of the unicellular excretory canal in *C. elegans*. Structure-function analysis of MRCK-1 indicates that the kinase domain, but not the small GTPase-binding CRIB domain, is required for canal outgrowth. Expression of a phosphomimetic form of MLC-4 rescues canal truncations in *mrck-1* mutants and shows enrichment at the growing canal tip. Specific depletion of MRCK-1 or MLC-4 in the canal causes severe truncations, with unlumenized projections of the basal membrane. Moreover, our work reveals a novel function for non-muscle myosin downstream of MRCK-1 in excretory canal outgrowth that may be conserved in the development of seamless tubes in other organisms.

## Introduction

Multicellularity necessitates the transport of liquids and gases throughout the organism, and as such the formation and maintenance of biological tubes is an essential process for metazoans and many other multicellular organisms. Large tubes are comprised of multiple cells surrounding an extracellular lumen, while the smallest tubes are comprised of a single cell with the lumen contained within the cytoplasm. Depending on their mechanism of formation, unicellular tubes may have an autocellular junction (‘seamed’ tubes), or completely lack tight and adherens junctions (‘seamless’ tubes). Whether they are multicellular or unicellular, biological tubes are formed through evolutionarily conserved biological processes: extracellular guidance cues, definition of apical-basal polarity, vesicle trafficking, and cytoskeletal remodeling (Herbert and Stainier, 2011; Lubarsky and Krasnow, 2003; Pradhan et al., 2022; Sundaram and Cohen, 2017; Weinstein, 2005; Xu and Cleaver, 2011).These processes must work in concert to transform single or multiple cells into a functional tube, and their dysregulation can cause aberrant tube formation.

Due to their simplicity, seamless tubes provide an ideal model to interrogate the conserved mechanisms of biological tube formation. Seamless tubes are found throughout the animal kingdom, where they compose portions of vertebrate microvasculature (Bär et al., 1984; Kamei et al., 2006; Wolff and Bär, 1972), as well as parts of organs such as the tip cells of the *Drosophila* trachea (Samakovlis et al., 1996), and the *C. elegans* excretory system (Kenneth Nelson et al., 1983). The largest tube in the excretory system of *C. elegans*, called the excretory canal cell, has been extensively studied to understand the molecular mechanisms of tube formation. The excretory canal is a seamless tube that maintains osmotic balance in the worm (Nelson and Riddle, 1984). This H-shaped cell has its cell body beneath the posterior bulb of the pharynx and extends two projections anteriorly and posteriorly along the sides of the body to span its entire length (Figure 1 A). These projections, called canals, are lumenized and connect to the central luminal space in the cell body, which then connects to the adjacent duct cell (Kenneth Nelson et al., 1983). The excretory canal cell is born prior to ventral enclosure during embryogenesis, and by the 1.5-fold stage the lumen of the canal has formed (Stone et al., 2009; Sulston et al., 1983). From this stage in embryogenesis until the end of the first larval stage the canal undergoes outgrowth as the anterior and posterior branches extend from the cell body and follow molecular guidance cues to reach their target positions at the distal ends of the worm (Fujita et al., 2003). The process of canal outgrowth is directed by many of the same attractant and repulsive guidance cues that direct neuronal outgrowth, and the leading tip of the canals has been described as analogous to the growth cone of an axon (Katidou et al., 2013; Kolotuev et al., 2013; Marcus-Gueret et al., 2012; McShea et al., 2013; Schmidt et al., 2009; Shaye and Greenwald, 2015; Stringham et al., 2002). For the remainder of development, the canal is in maintenance phase, growing at the same rate as the rest of the body to maintain its relative position.

**Figure 1:**
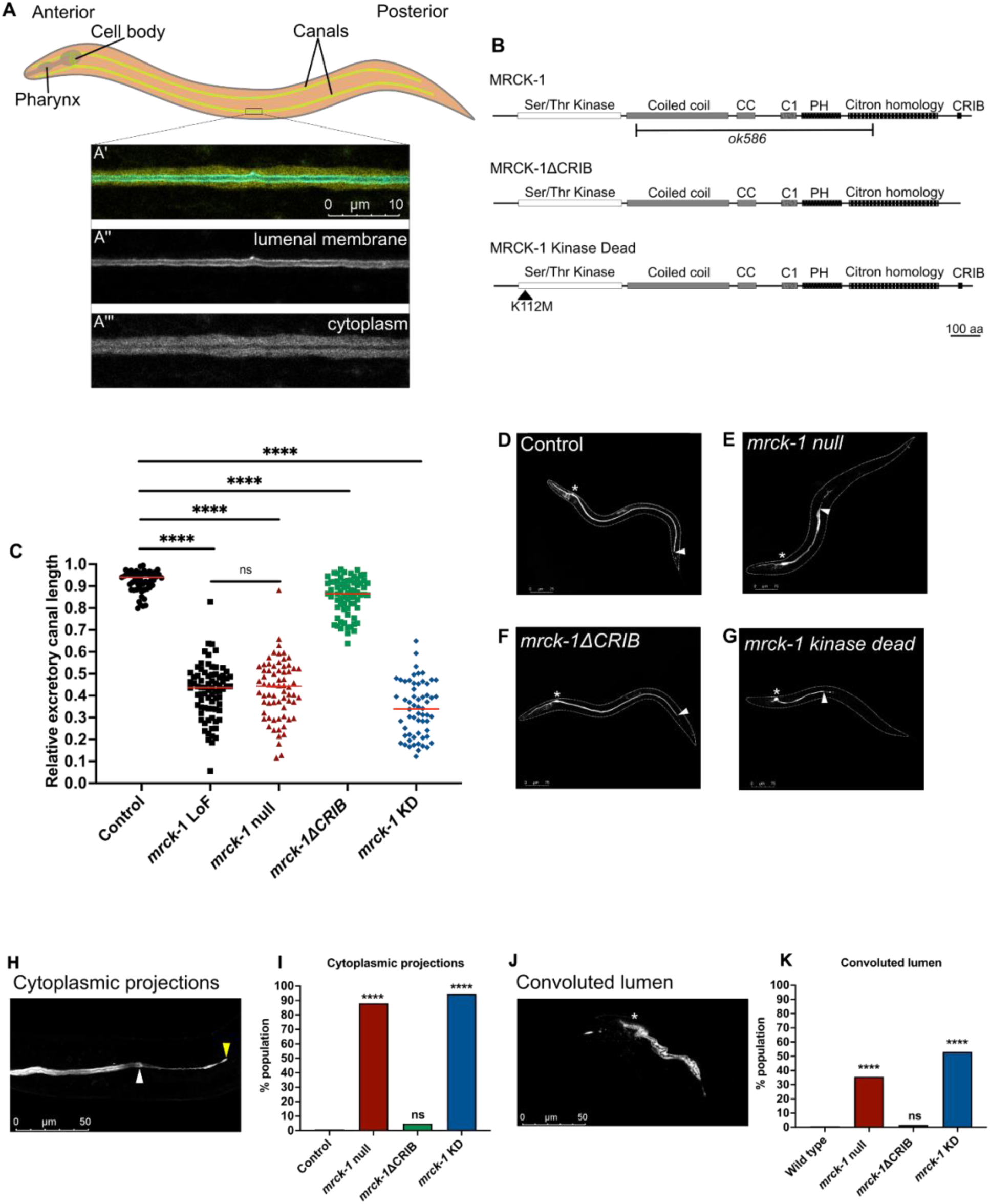
Loss of *mrck-1* or mutation of its kinase domain causes severe truncations and morphological defects in the excretory canal. **(A-A’’’)** The excretory cell of *C. elegans* is an H-shaped unicellular tube that extends canals anteriorly and posteriorly along the length of the body. **(A’)** This microscopy image of a section of the canal in a wild type worm highlights the apical/luminal membrane **(A’’)** (cyan, canalp::ifb-1::CFP) and the cytoplasm **(A’’’)** (yellow, canalp::YFP). **(B)** Structure of wild type MRCK-1 and the predicted protein products of *mrck-1ΔCRIB* and *mrck-1* kinase dead (KD) alleles. *ok586* shows the *mrck-1(ok586)* deletion allele in reference to the wild type MRCK-1 amino acid sequence, although the effect of the allele on the protein has not been experimentally determined. This 2447 bp deletion/124bp insertion is predicted to disrupt splice sites and cause missense mutations. CC, coil coiled domain; C1, protein kinase C conserved domain 1; PH, Pleckstrin homology; CRIB, Cdc42 and Rac-interactive binding. (C-G) *mrck-1* LoF (n=77), *mrck-1* null (n=68) and KD mutants (n=60) have severe canal truncations, while *mrck-1ΔCRIB* (n=71) mutants have modest truncations. Red lines in the graph denote the medians, **** p < 0.0001 versus wild type control (n=58) (Mann-Whitney test). **(H-K)** A significant proportion of *mrck-1* null and KD mutants show a failure of the lumen to extend to the end of the canal (cytoplasmic projections) and a folding up of the lumen within the cytoplasm of the cell (convoluted lumen). n ≥ 100 for each genotype (see Materials for details). ns p ≥ 0.05, **** p < 0.0001 versus wild type control (Fisher’s exact test). Asterisks denote the location of the cell body of the excretory cell, white arrowhead denotes the end of the lumenized portion of the canal, yellow arrowhead denotes the end of the canal cytoplasm.

During outgrowth and maintenance, the canal requires definition of apical-basal polarity partnered with extensive remodeling of the cytoskeleton and dynamic vesicle trafficking. The apical (luminal) and basal domains of the tube are defined by the PAR polarity proteins for polarized fusion of vesicles to the correct membrane (Abrams and Nance, 2021; Armenti et al., 2014a). Small GTPases including RAL-1 and CDC-42 are essential regulators of trafficking and polarity in the excretory canal through regulation of the PAR proteins and exocyst complex (Abrams and Nance, 2021; Armenti et al., 2014a; Lant et al., 2015; Mattingly and Buechner, 2011). Upstream regulators of vesicle trafficking, including CCM3/STRIPAK and the EXC-9/EXC-1/EXC-5(GEF) signaling cascade have also been identified as essential components of canal extension (Armenti et al., 2014a; Gao et al., 2001; Grussendorf et al., 2016; Lant et al., 2015; Suzuki et al., 2001; Tong and Buechner, 2008). Cytoskeletal components including actin, intermediate filaments, microtubules, and cytoskeletal linker proteins ERM-1 (ezrin-radixin-moesin) and SMA-1/β-H-spectrin are required for structural support and to regulate vesicle trafficking at the luminal membrane of the canal (Al-Hashimi et al., 2018; Fujita et al., 2003; Göbel et al., 2004; Khan et al., 2019, 2013; Kolotuev et al., 2013; Shaye and Greenwald, 2015; Woo et al., 2004).

Previously our lab identified the serine/threonine kinase *mrck-1* (myotonic dystrophy-related Cdc42-binding kinase homolog 1) as a regulator of canal extension downstream of the disease-associated gene *ccm-3,* but the role of MRCK-1 in the canal remains largely uncharacterized (Lant et al., 2015). The myotonic dystrophy-related Cdc42-binding kinases (MRCKs) are highly conserved in animals including humans, where there are three paralogs: MRCKɑ, MRCKβ, and MRCKƔ (Unbekandt and Olson, 2014). These kinases are part of the larger dystonia myotonica protein kinase (DMPK) family, all members of which promote the activation of the actomyosin complex through phosphorylation of the regulatory light chain of non-muscle myosin (Amano et al., 1996; Leung et al., 1998; Luo et al., 1997; Murányi et al., 2001; Yamashiro et al., 2003; Zhao et al., 1997). In vertebrate models the MRCKs regulate cell-cell junction remodeling, apical polarity, and cell migration through actomyosin (Ando et al., 2013; Gomes et al., 2005; Huo et al., 2011; Tan et al., 2011; Zihni et al., 2017) and MRCK has been shown to interact with the small GTPase CDC42 through its Cdc42 and Rac-interactive binding (CRIB) domain (Leung et al., 1998; Luo et al., 1997). MRCKꞵ has been implicated in lumen formation of endothelial cells in 3D culture downstream of Cdc42, but the mechanism by which it contributes to this process is unknown (Barry et al., 2016; Koh et al., 2008; Norden et al., 2016). In *C. elegans mrck-1* promotes the phosphorylation of the regulatory light chain homolog MLC-4 during embryonic elongation (Gally et al., 2009) and functions with active CDC-42 to activate non-muscle myosin during early gastrulation (Marston et al., 2016).

To expand upon our discovery of *mrck-1* as a component of seamless tube formation we aimed to define the role of MRCK-1 in excretory canal development and identify the downstream pathway(s) it regulates in this role. Here we show that MRCK-1 kinase domain but not its CRIB domain is required for canal outgrowth, and that maternally contributed *mrck-1*/MRCK-1 functions during this early stage of canal development. Mechanistically, we show that phosphorylated MLC-4 acts downstream of *mrck-1* to rescue canal defects in *mrck-1* mutants. Additionally, we demonstrate a requirement for MLC-4 during excretory canal growth, which phenocopies the canal defects caused by loss of MRCK-1. This work defines a novel role for MRCK-1/non-muscle myosin in seamless tube outgrowth, which may be conserved in other contexts including vertebrate tubulogenesis.

## Results

### MRCK-1 kinase domain is required for excretory canal extension

Given the established role of the MRCKs as kinases and Cdc42-binding proteins (Leung et al., 1998; Luo et al., 1997) we tested the requirement of the kinase and Cdc42 and Rac-interactive binding (CRIB) domains for MRCK-1 function in the excretory canal. Using CRISPR/Cas9 gene editing we edited *mrck-1* at its endogenous locus to create a CRIB domain deletion allele (*mrck-1ΔCRIB*) that removes the last 58 amino acids of the protein, which includes the 14 amino acid CRIB domain (Figure 1 B). We created a kinase dead (KD) allele that encodes a K112M amino acid substitution (Figure 1 B), which is predicted to abrogate kinase activity by disrupting ATP-binding (Sumi et al., 2001; Wilkinson et al., 2005). We also created an *mrck-1* null allele by deleting the entire coding sequence of the gene (Figure 1 B), as a positive control for complete loss of MRCK-1. Additionally we analyzed a published *mrck-1* loss of function allele (*ok586*) which we used previously to test the requirement of *mrck-1* for canal extension (Lant et al., 2015; The *C. elegans* Deletion Mutant Consortium, 2012). The *mrck-1* LoF allele encodes a complex substitution (2447 bp deletion and 124 bp insertion) predicted to cause splice site variants and missense mutations in the protein which abolish its function.

While excretory canals of wild type worms extend nearly to the end of the worm (median 0.94 relative length), we found that the *mrck-1* null and KD mutants had severely truncated canals that were less than half the worm’s body length (median 0.44 and 0.34) (Figure 1 C, E, G). The *mrck-1ΔCRIB* mutants also had canals that were significantly shorter than wild type, but the level of truncations was mild compared to the other *mrck-1* mutants (median 0.87) (Figure 1 C, F). We found that there is no significant difference between the canal lengths of *mrck-1* LoF and *mrck-1* null mutants, so we conclude that the LoF allele causes complete loss of *mrck-1* function (Figure 1 C).

The canals of *mrck-1* null and KD mutants show two other morphological defects that were absent in wild type and *mrck-1ΔCRIB* mutant worms: cytoplasmic projections and convoluted lumens. Cytoplasmic projection is a defect in which the basal membrane and cytoplasm of the canal extend without the luminal membrane, creating an unlumenized section at the tip of the canal (Figure 1 H) (Khan et al., 2013; Lant et al., 2015). Cytoplasmic projections were observed in less than 1% of wild type worms and only 6.7% of *mrck-1ΔCRIB* mutants but occurred in the majority of *mrck-1* null and KD mutant populations (88.7 % and 95.0 %) (Figure 1 I). Convoluted lumen describes defects in which the lumen of the canal appears ‘folded’ up in the cytoplasm of the cell (Figure 1 J) (Buechner, 2002; Buechner et al., 1999), which was observed in less than 2% of wild type worms and *mrck-1ΔCRIB* mutants, but was frequent in *mrck-1* null and KD mutants (37.1 % and 53.4 %) (Figure 1 K).

Overall, the similarities in canal truncations and other morphological defects between the *mrck-1* null and KD mutants suggests that the kinase domain is essential for MRCK-1 function in the canal. The mild canal truncations and rare morphological defects in the *mrck-1ΔCRIB* mutants suggests that this domain, and potentially an interaction with CDC-42, is not necessary for MRCK-1 function in canal extension.

### *mrck-1* is required for canal outgrowth

The previous evaluation of canal lengths in *mrck-1* mutants was performed on late L4 stage and early adult worms when the canal has fully developed. To gain insight into the timing of *mrck-1* function during development we next investigated when these truncations first appeared. The development of the excretory canal can be divided into two discrete phases: outgrowth and maintenance (Figure 2 A). Starting just after the lumen is formed at 1.5-fold stage of embryogenesis until the end of the first larval stage the canal undergoes outgrowth where the anterior and posterior tips follow molecular guidance cues that extend the canals to reach their target positions (Figure 2 A). After this, the canal switches to a maintenance phase where it grows at the same rate as the body to maintain its relative position (Figure 2 A) (Buechner, 2002; Fujita et al., 2003).

**Figure 2:**
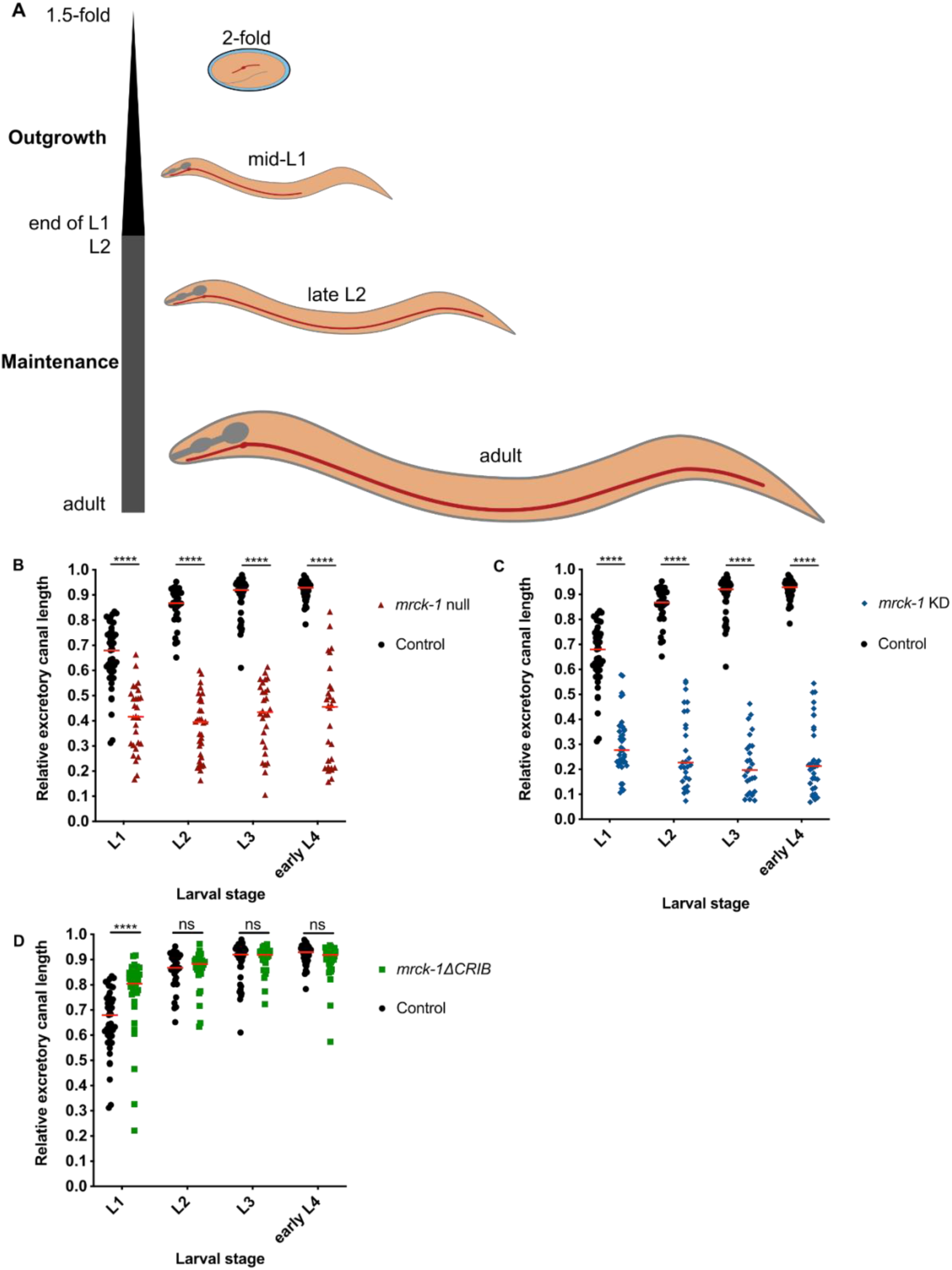
*mrck-1* and its kinase domain are required for excretory canal outgrowth. **(A)** From the 1.5-fold stage during embryogenesis to the end of the first larval stage (L1) the excretory canal undergoes outgrowth, where it follows guidance cues to reach its final position near the tail of the worm. For the rest of development, the canal undergoes maintenance, where it grows at the same rate at the body to maintain its relative position. Worms are not to scale. The *mrck-1* null **(B)** and KD **(C)** mutants have severe canal truncations by L1, and these defects persist throughout development. (**D)** *mrck-1ΔCRIB* mutants do not have canals shorter than wild type controls at any stage of larval development. Sample size (L1, L2, L3, eL4): *mrck-1* null n=28, 32, 28, 28; *mrck-1* KD n=39, 27, 26, 32; *mrck-1ΔCRIB* n=34, 29, 27, 35; control n=44, 30, 39, 33. Red lines in the graphs denote the medians. ns p ≥ 0.05, **** p < 0.0001, versus wild type control (Mann-Whitney test).

We compared the relative canal lengths of *mrck-1* null, KD, and *mrck-1ΔCRIB* mutants to wild type worms at each larval stage to determine when truncations first appear (Figure 2 B-D). The *mrck-1* null and KD mutants have significantly shorter canals than wild type worms by the first larval stage, and these truncations persist throughout development (Figure 2 B, C). This suggests that *mrck-1* kinase activity is required during the outgrowth phase of canal development. *mrck-1ΔCRIB* mutants do not have shorter canals than wild type worms at any larval stage, which suggests that an interaction with CDC-42 is not required for MRCK-1 function during canal outgrowth and much of the maintenance phase (excluding late L4/early adult) (Figure 2 D).

### Maternally supplied *mrck-1* promotes canal extension

When quantifying the canal truncations of the *mrck-1* null and KD adults, we observed that the canals of the KD mutants are significantly shorter than the null mutants (Figure 3 A). Initially we hypothesized the *mrck-1* KD allele may be acting in a dominant negative fashion, but worms heterozygous for the allele have wild type canals (Figure S1).

**Figure 3:**
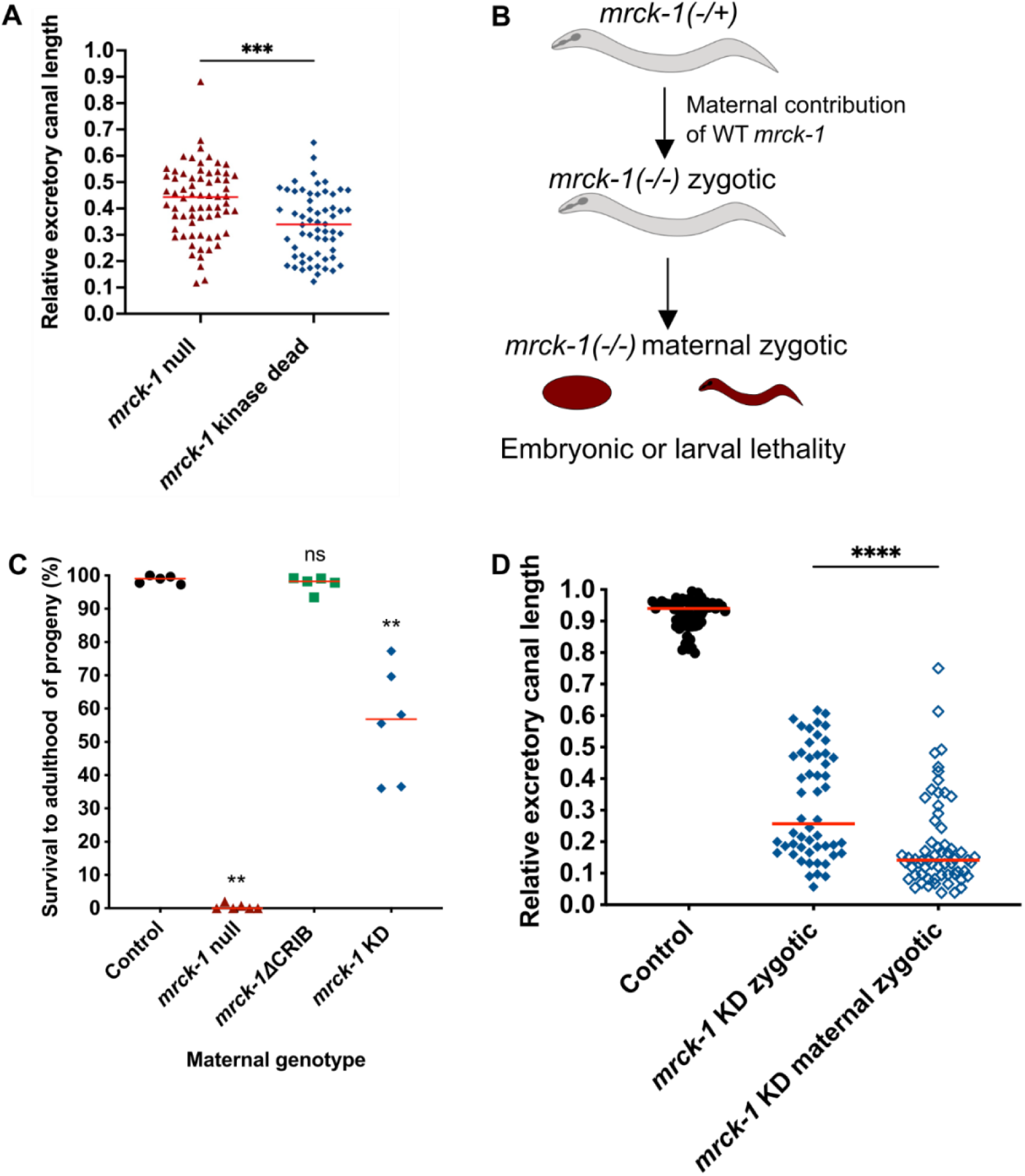
Maternally supplied *mrck-1*/MRCK-1 suppresses canal truncations in *mrck-1* KD mutants. **(A)** *mrck-1* KD mutants have significantly shorter canals than *mrck-1* null mutants. *** p < 0.001 (Mann-Whitney test). **(B)** *mrck-1* zygotic mutants survive embryonic elongation due to maternally contributed wild type *mrck-1*/MRCK-1, while *mrck-1* maternal zygotic mutants die as embryos or L1s. **(C)** While the survival of *mrck-1* null mutant progeny is less than 1%, a median of 57% of *mrck-1* KD mutant progeny survive to adulthood. ns p ≥ 0.05, ** p < 0.01, versus wild type control (Mann-Whitney test). **(D)** The canals of *mrck-1* KD maternal zygotic mutants (n=52) are significantly shorter than the zygotic mutants (n=62). *** p < 0.001 (Mann-Whitney test). Red lines in the graphs denote the medians.

Next, we investigated the possibility that the less severe truncations observed in *mrck-1* null mutants may be due to maternal rescue. In addition to its function in the canal, *mrck-1* is required during embryogenesis for the activation of non-muscle myosin during embryonic elongation (Gally et al., 2009). Zygotic (Z) *mrck-1* mutants from heterozygous mothers survive to adulthood due to maternally contributed wild type *mrck-1*, while the maternal zygotic (MZ) *mrck-1* mutants die during embryogenesis or at the first larval stage (Figure 3 B). We measured the maternal embryonic/larval lethal phenotype of the *mrck-1* alleles we generated as the survival to adulthood of progeny from Z mutant mothers (as a percent of total progeny laid) and compared this to wild type controls (Figure 3 C). Almost all progeny from wild type control mothers survive to adulthood, compared to less than 1% of progeny from *mrck-1* null mutants (Figure 3 C). The *mrck-1* KD mutants showed an intermediate phenotype where a median of 56.8 % of progeny survived to adulthood (Figure 3 C). Given these results, it’s likely that the original population of *mrck-1* KD mutants was a combination of Z and MZ mutants, which could account for the more severe canal truncation phenotype. In agreement with this hypothesis, when we compared the canal lengths of Z and MZ *mrck-1* KD mutants we found that the MZ mutants have significantly shorter canals (Figure 3 D). These findings suggest that maternally contributed *mrck-1* plays a role in excretory canal development and indicate that the effect of total loss of *mrck-1* is partially suppressed in the previously quantified Z *mrck-1* null mutants.

### MRCK-1 functions autonomously in the canal for outgrowth

To circumvent the maternal contribution of *mrck-1*, we used the tissue-specific ZIF-1/ZF1 protein degradation system, which was previously adapted to degrade target proteins rapidly and specifically in the canal (Abrams and Nance, 2021). This system uses tissue-specific expression of the *C. elegans* substrate-recognition protein ZIF-1 to recruit target proteins tagged with a ZF1 recognition motif to the E3/E2 ubiquitin ligase complexes for poly-ubiquitination and subsequent proteasome-mediated degradation (Armenti et al., 2014b, 2014a). To create the canal-specific MRCK-1 ablation (MRCK-1^canal-^) mutants, we added the ZF1 recognition motif sequence to the C-terminus of *mrck-1* at its endogenous locus in a strain with a transgene that expresses ZIF-1 under a canal-specific promoter. These MRCK-1^canal-^ mutants had canals that were significantly shorter than wild type, demonstrating an autonomous function for MRCK-1 in the canal (Figure 4 A-C). MRCK-1^canal-^ mutants also have significantly shorter canals than the previously quantified *mrck-1* null mutants, providing further evidence that maternally contributed *mrck-1* functions in canal extension (Figure S2).

**Figure 4:**
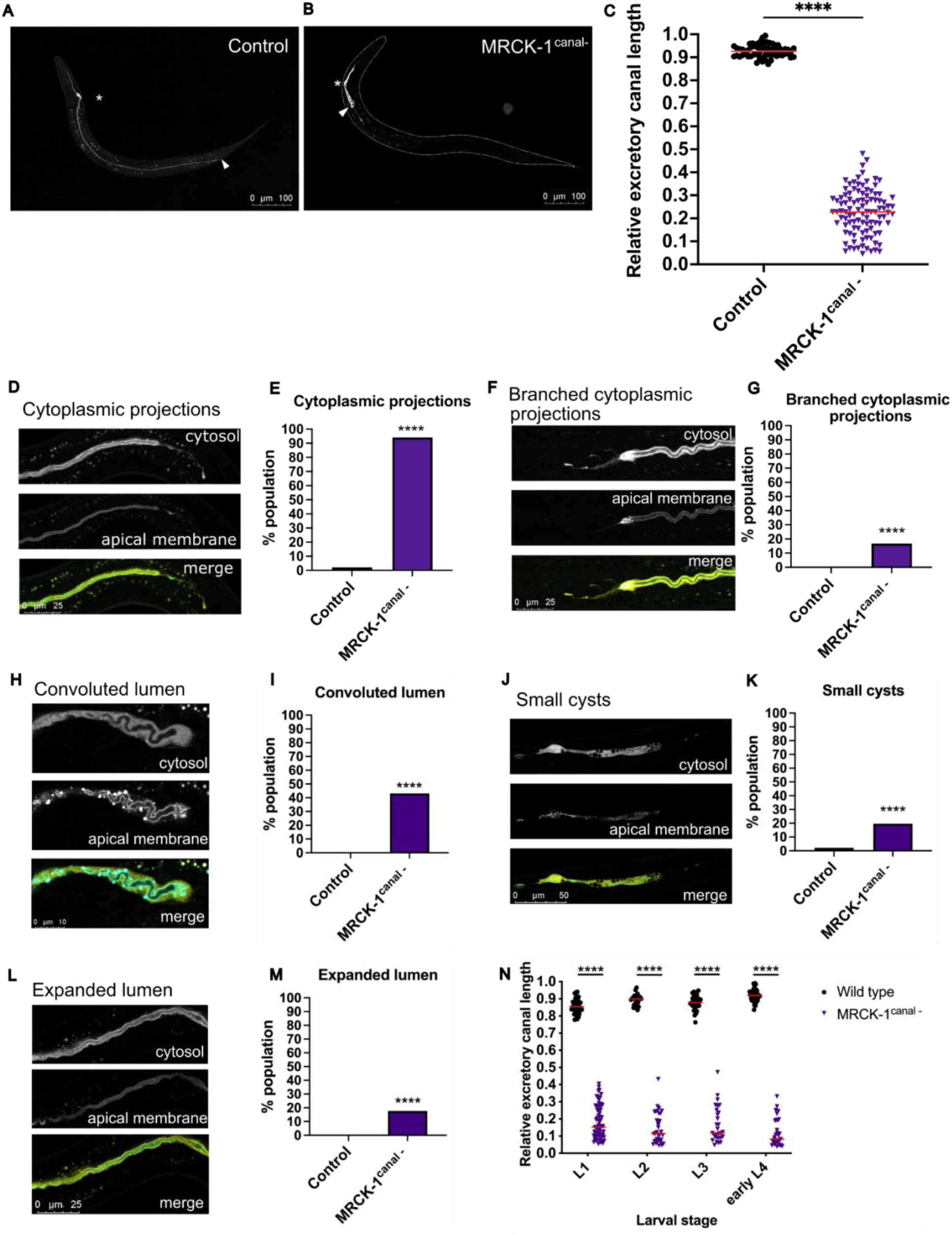
MRCK-1 functions autonomously in the excretory canal cell during outgrowth. **(A-C)** MRCK-1^canal-^ adults (n=102) with MRCK-1 ablated only in the excretory canal cell have extremely truncated canals that are significantly shorter than wild type controls (n=95). **** p < 0.0001 (Mann-Whitney test). Asterisks denote the location of the cell body of the excretory cell, white arrowhead denotes the end of the lumenized portion of the canal. MRCK-1^canal-^ mutants have several morphological defects including **(D)** cytoplasmic projections, **(F)** branched cytoplasmic projections, **(H)** convoluted lumens, **(J)** small cysts, and **(L)** expanded lumens. All these defects were observed at a significantly higher frequency in MRCK-1^canal-^ mutants compared to wild type controls **(E, G, I, K, M)**. **** p < 0.0001 (Fisher’s exact test), n ≥ 95 (see Materials for details). **(N)** By the first larval stage the canals of MRCK-1^canal-^ mutants are significantly shorter than wild type controls, and these truncations persist throughout development. Sample size (L1, L2, L3, eL4): MRCK-1^canal-^ n=66, 33, 25, 19; control n=34, 28, 30, 44. **** p < 0.0001 (Mann-Whitney test). Red lines on graphs denote population medians.

MRCK-1^canal-^ mutants express a fluorescent reporter that marks the luminal membrane of the canal with CFP and the cytosol with YFP, which allowed for identification of morphological defects that were not apparent with the previously used cytosol-only canal marker. In addition to cytoplasmic projections and convoluted lumen defects, we identified 3 additional morphological defects in MRCK-1^canal-^ mutants: branched cytoplasmic projections, small cysts, and expanded lumen (Figure 4 F, J, L). Branched cytoplasmic projections are a subtype of the cytoplasmic projections defect, where the projection at the tip of the canal becomes two or more separate branches (Figure 4 F). Small cysts are spherical structures smaller than the diameter of the canal contained in the cytoplasm of the cell (Figure 4 J). The expanded lumen phenotype describes canals in which most of the excretory canal volume is taken up by lumen (Figure 4 L). All these defects were observed at a significantly higher frequency in MRCK-1^canal-^ mutants than wild type controls, where they seldom or never were observed (Figure 4 E, G, I, K, M).

After analyzing the canals of adult MRCK-1^canal-^ mutants, we were interested in characterizing when these defects arise during development. Similar to the *mrck-1* null and KD mutants, MRCK-1^canal-^ mutants have canals that are significantly shorter than wild type beginning at the first larval stage and persisting into adulthood (Figure 4 N). There was a significant increase in the frequency of expanded and convoluted lumens in the MRCK-1^canal-^ mutants throughout development (Figure S3 D, E) while there is no significant difference in the frequency of cytoplasmic projections (single or branched), or small cysts (Figure S3 A-C). This suggests that the cytoplasmic projections and cyst defects are a consequence of loss of MRCK-1 during canal outgrowth. The expanded and convoluted lumens defects may indicate a requirement for MRCK-1 during the maintenance phase or could be a general consequence of the lumen attempting to grow in truncated canals.

### Expression of a phosphomimetic MLC-4 mutant rescues canal truncations in *mrck-1* mutants

Since the requirement of the kinase domain supports a function for MRCK-1 kinase activity in the canal, we next wanted to identify genes which could be downstream kinase substrates. We hypothesized that MRCK-1 may function through its known target the regulatory light chain (RLC) of non-muscle myosin (MLC-4). MRCK-1 is known to promote phosphorylation of MLC-4 during early gastrulation and embryonic elongation to generate contractile force (Gally et al., 2009; Marston et al., 2016). Non-muscle myosin is activated by phosphorylation of two conserved residues on the RLCs, threonine 17 and serine 18 in MLC-4 (Gally et al., 2009). To test a role for *mlc-4* downstream of *mrck-1* in canal extension, we replaced threonine 17 and serine 18 with aspartic acids (MLC-4DD) to generate a phosphomimetic protein, which was expressed in *mrck-1* LoF mutants. Expression of the MLC-4DD phosphomimetic mutant was able to rescue canal truncations in the *mrck-1* mutants, which suggests that phosphorylation of *mlc-4* functions downstream of *mrck-1* for canal extension (Figure 5 A, B). MRCKs have been shown to phosphorylate the RLC of non-muscle myosin directly (Leung et al., 1998), or through inhibitory phosphorylation of myosin light chain phosphatase (MLCP), which is encoded by *mel-11* in *C. elegans* (Gally et al., 2009; Tan et al., 2001; Wilkinson et al., 2005). To determine if *mel-11* acts downstream of *mrck-1* in the canal, we knocked down its expression in *mrck-1* null mutants (Figure S4). Knockdown of *mel-11* did not rescue canal truncations in the *mrck-1* mutants (Figure S4), which suggests that *mrck-1* does not inhibit this phosphatase to promote phosphorylation of MLC-4 for canal extension.

**Figure 5:**
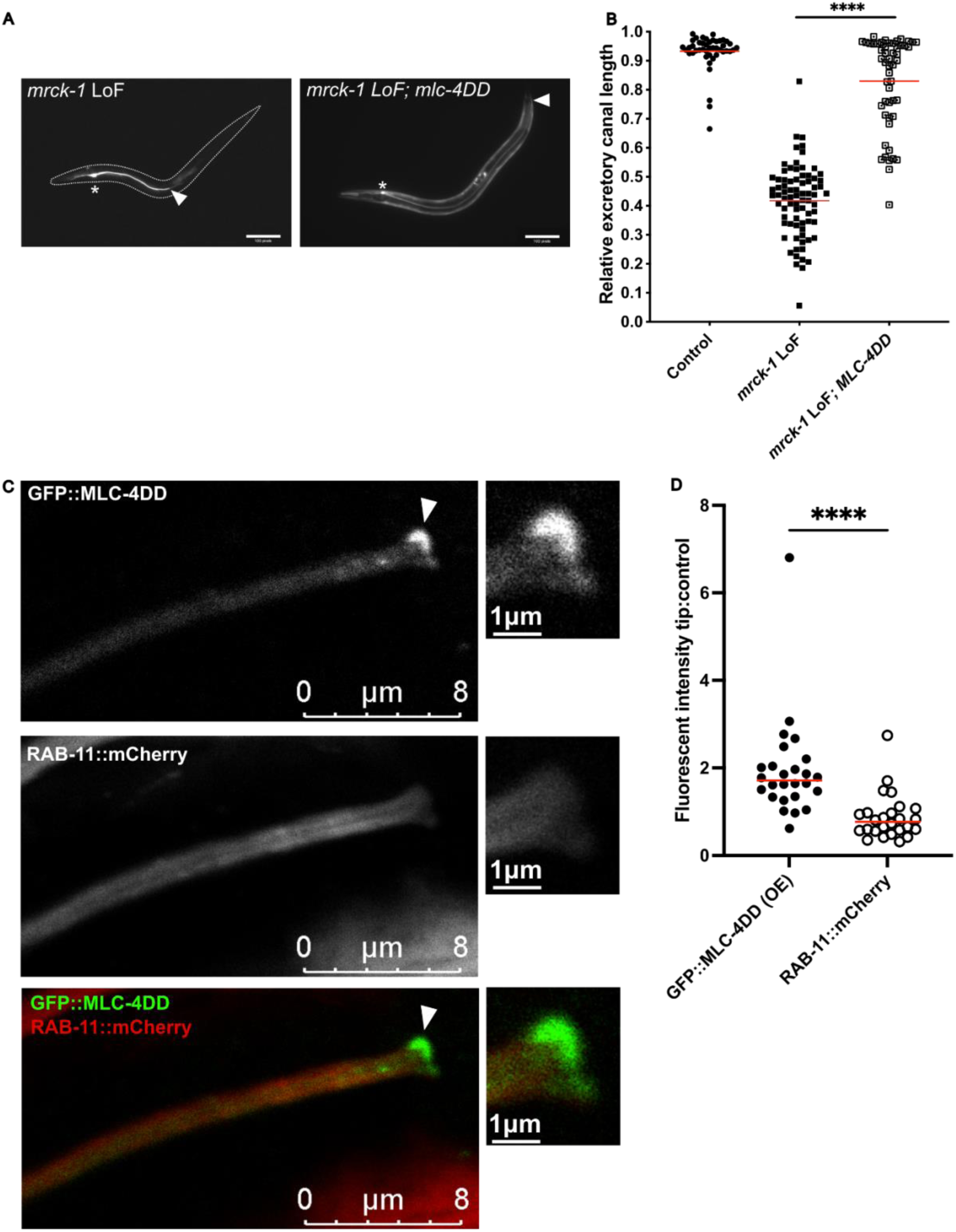
Expression of an MLC-4 phosphomimetic mutant rescues canal truncations in *mrck-1* mutants. **(A)** Expression of *mlc-4DD* in the canal of *mrck-1* LoF mutants can restore canals to wild type lengths. **(B)** *mrck-1* mutants with the *mlc-4DD* transgene (n=52) have significantly longer canals than the single mutants (n=77). Wild type control n=46. **** p < 0.0001 (Mann-Whitney test). Red lines on graphs denote population medians. **(C)** GFP::MLC-4DD expressed in the canal shows enrichment at the growing tip of the canal during outgrowth in L1 larva (white arrow). This enrichment at the growing tip was not observed in the control RAB-11::mCherry marker, which has a diffuse cytoplasmic localization. **(D)** L1 worms expressing *gfp::mlc-4DD* in the canal show enrichment of GFP::MLC-4DD at the tip of the canal, compared to an adjacent control region. RAB-11::mCherry fluorescence is slightly diminished at the canal tip compared to the control region. n=26. **** p < 0.0001 (paired samples Wilcoxon test).

To better understand the function of phosphorylated MLC-4 in the excretory canal, we observed the localization of the *mlc-4* phosphomimetic mutant tagged with GFP (GFP::MLC-4DD) expressed under a canal-specific promoter. Localization of this translational reporter was compared with a RAB-11::mCherry reporter expressed in the canal, which shows diffuse localization throughout the cytoplasm. We found that in L1s, when the canal is in outgrowth stage, GFP::MLC-4DD is enriched at the growing tip of the canal (Figure 5 C, D). This enrichment at the growing tip is unique to GFP::MLC-4DD, whereas the RAB-11::mCherry marker is not enriched at the canal tip (Figure 5 C, D).

Activated non-muscle myosin binds to F-actin through the head domain of the heavy chain to form actomyosin, the protein complex that exerts contractile force upon structures within the cell. Given the interaction between actin and non-muscle myosin, and the enrichment of F-actin (Shaye and Greenwald, 2015) and GFP::MLC-4DD at the tip of the canal, we wanted to see if F-actin organization was perturbed upon loss of *mrck-1*. Localization of F-actin was carried out using a LifeAct::TagRFP reporter expressed in the excretory canal. At L1 stage during canal outgrowth the LifeAct::TagRFP reporter showed enrichment growing tip, demarking the previously described leading edge F-actin structure (Shaye and Greenwald, 2015) (Figure S5 A). This localization of F-actin at the leading edge did not change in the *mrck-1* mutants (Sup Figure S5), indicating that F-actin organization is not detectably affected by this kinase in the canal.

### MLC-4 functions autonomously to promote canal outgrowth

Since our genetic evidence suggests that *mlc-4* functions downstream of *mrck-1* in the canal, we wanted to determine if *mlc-4* functioned autonomously to promote canal outgrowth. *mlc-4* is an essential gene due to its roles in cytokinesis and polarity during early embryogenesis (Shelton et al., 1999) and elongation later in embryonic development (Gally et al., 2009). Therefore, to avoid lethality from complete loss of *mlc-4*, we turned to the ZIF-1/ZF1 protein degradation system again to specifically ablate MLC-4 in the canal.

We found that MLC-4^canal-^ mutants had severe canal truncations supporting an autonomous role for MLC-4 in promoting canal extension (Figure 6 A, B). Notably the canal lengths of MLC-4^canal-^ and MRCK-1^canal-^ mutants are not significantly different, which is consistent with our observations that MLC-4DD rescues canal truncations in *mrck-1* mutants (Figure 5 B). Like MRCK-1^canal-^ mutants, the canals of MLC-4^canal-^ mutants have single and branched cytoplasmic projections, small cysts, convoluted lumens, and expanded lumens (Figure 6 C-G). Although MLC-4^canal-^ and MRCK-1^canal-^ mutants have the same kinds of morphological defects, some of these defects were present at different frequencies. There was no significant difference in the frequency of cytoplasmic projections defects, as they were present in almost all MLC-4^canal-^ and MRCK-1^canal-^ mutants (Figure 6 C). The branched cytoplasmic projections and expanded lumens were approximately twice as prevalent in MLC-4^canal-^ mutants compared to MRCK-1^canal-^ mutants, while small cysts were three times as prevalent (Figure 6 D, F, G). Conversely, convoluted lumens were twice as prevalent in MRCK-1^canal-^ mutants (Figure 6 E).

**Figure 6:**
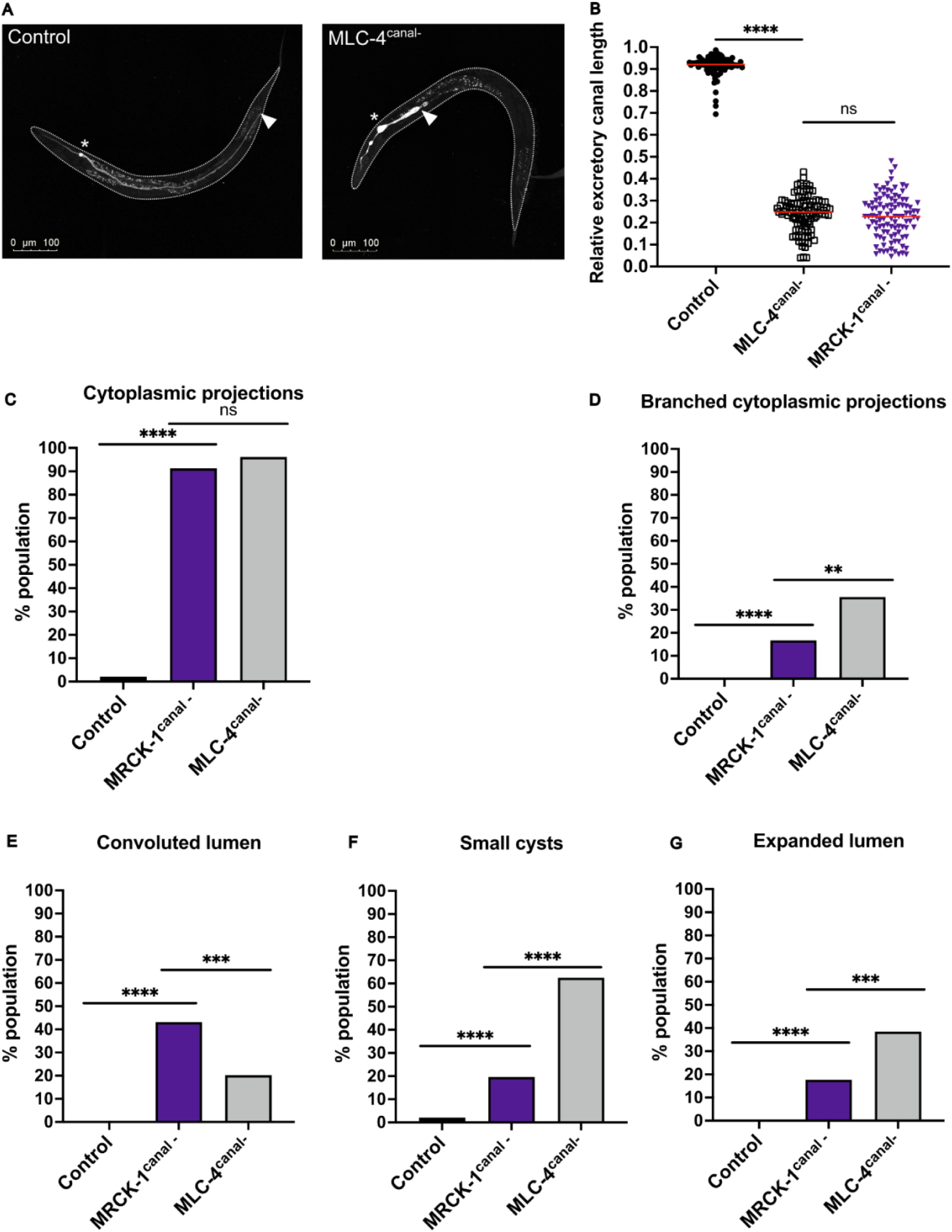
Ablation of MLC-4 in the excretory cell causes canal truncations and morphological defects that resemble MRCK-1 ablation. **(A-B)** MLC-4^canal-^ mutants (n=104) have significantly shorter canals than wild type controls (n=91), but there is no significant difference between these mutants and the canals of MRCK^canal-^ mutants (n=102). ns p ≥ 0.05, **** p < 0.0001 (Mann-Whitney test). Red lines on graphs denote population medians. **(C-G)** MLC-4^canal-^ mutants display the same kinds of canal defects as MRCK-1^canal-^ mutants. **(C)** Cytoplasmic projections are present at equal frequencies in MLC-4^canal-^ and MRCK^canal-^ mutants, but the other defects are more frequent in one mutant. **(D, F, G)** Branched cytoplasmic projections, small cysts, and expanded lumens are more prevalent in MLC-4^canal-^ mutants, while **(E)** convoluted lumens are more prevalent in MRCK^canal-^ mutants. MLC-4^canal-^ (n=104), MRCK^canal-^ (n=102), wild type controls (n=91). ns p ≥ 0.05, ** p < 0.01, *** p < 0.001, **** p < 0.0001 (Fisher’s exact test).

Canal truncations in MLC-4^canal-^ mutants were observed in the first larval stage during canal outgrowth and persisted into adulthood (Figure 7), which suggests a role for non-muscle myosin in canal outgrowth. Similar to MRCK-1^canal-^ mutants, Cytoplasmic projections were observed at a high frequency in MLC-4^canal-^ mutants starting at L1 stage and did not significantly change during development (Figure S6), which suggests that this defect arises during outgrowth. The frequency of expanded and convoluted lumens was very low to absent in the MLC-4^canal-^ mutants at L1 stage, and significantly increased throughout development to adulthood (Figure S6). This pattern is similar to the development expanded and convoluted lumens in the MRCK-1^canal-^ mutants, and could indicate a requirement for MLC-4 later in development, or a general consequence of extremely truncated canals. Unlike the MRCK-1^canal-^ mutants, the frequency of small cysts significantly increased in the MLC-4^canal-^ mutants after the 2^nd^ larval stage (Fig S6), which could indicate a role for non-muscle myosin in trafficking during maintenance phase in addition to a role during outgrowth.

**Figure 7:**
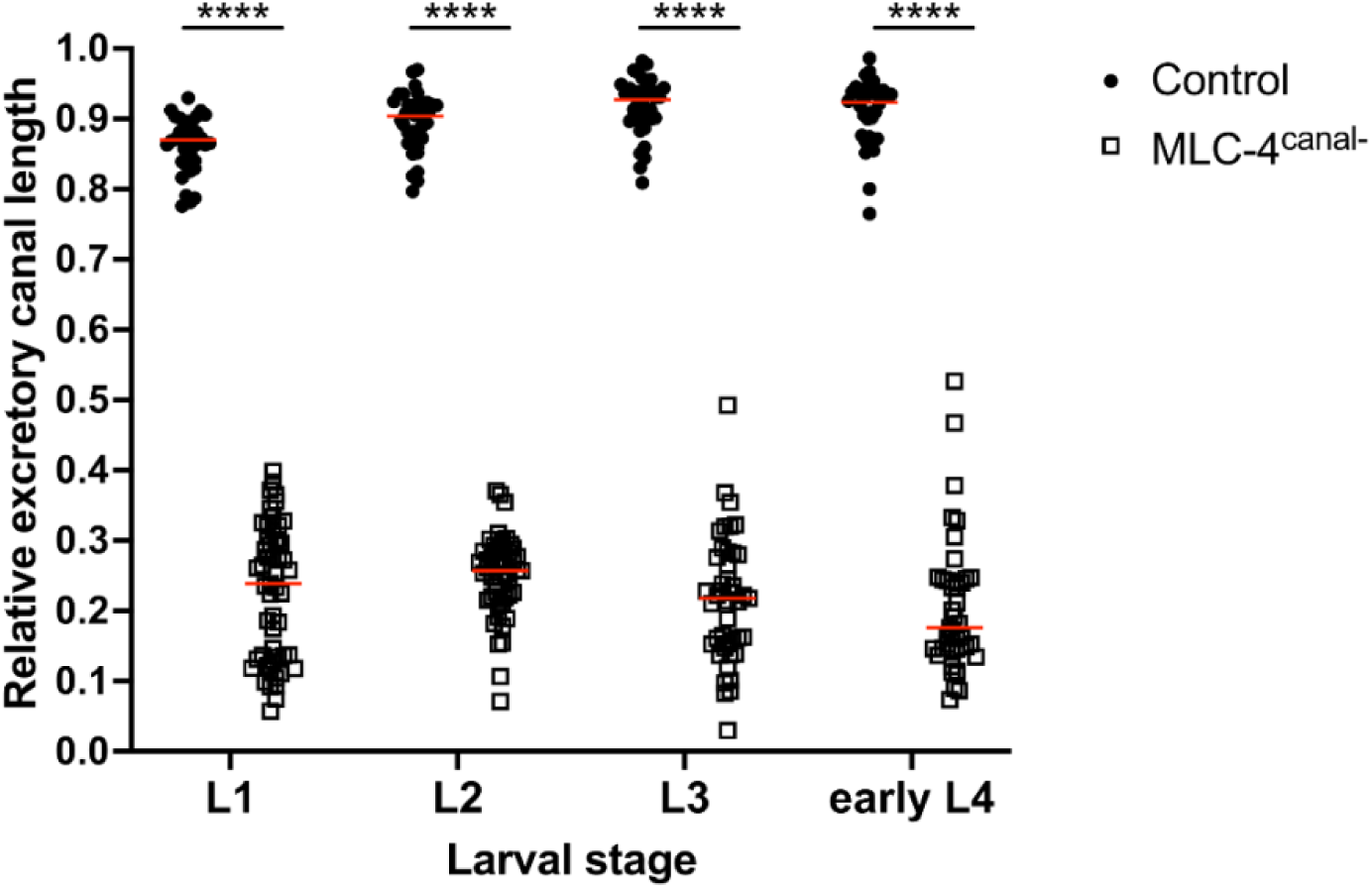
MLC-4 is required in the excretory canal during outgrowth. By L1 stage the canals of MLC-4^canal-^ mutants are significantly shorter than wild type controls, and these truncations persist throughout development to adulthood. Sample size (L1, L2, L3, eL4): MLC-4^canal-^ n=53, 51, 43, 43; wild type control n=38, 36, 38, 36. **** p < 0.0001 (Mann-Whitney test). Red lines on graphs denote population medians.

## Discussion

The formation and patterning of tubes is an essential process for many organs and structures in metazoans, and the molecular mechanisms that govern these processes are highly conserved. Here, we have defined an autonomous role of the conserved kinase MRCK-1 in the development of the seamless excretory canal cell in *C. elegans.* Mutations that disrupt the kinase domain, but not the Cdc42-binding CRIB domain, prevent outgrowth of the canal. Consistent with these observations, previous work showed that Cdc42 is not required for MRCKA kinase activity (Leung et al., 1998), although most previously characterized functions of MRCK kinases *in vivo* or *in vitro* require CDC42-dependent localization at specific cell membrane sites for its proper function (Ando et al., 2013; Huo et al., 2011; Marston et al., 2016; Zihni et al., 2017). In the *Drosophila* visual system the MRCK homolog *gek* is required for axon targeting in a CRIB-independent manner, although the phenotypic similarities between *gek* and *cdc42* mutants led the authors to conclude that this is still a Cdc42-dependent process (Gontang et al., 2011). Ablation of CDC-42 in the excretory canal has been reported to cause less severe truncations that occur later in development than what we observe in our *mrck-1*/MRCK-1^canal-^ mutants, and these defects are caused through a loss of polarized localization of the exocyst complex (Abrams and Nance, 2021). Previously we proposed that MRCK-1 functions downstream of *cdc-42* to promote endocytic recycling in the canal (Lant et al., 2015), but our findings suggest that MRCK-1 functions independently of CDC-42 in this context. In addition to the CRIB domain, localization of MRCK-1 to the cell membrane is mediated by a tripartite module of the C1, PH, and CNH domains, which bind preferentially to specific phosphatidylinositol phosphates (PIPs) (Truebestein et al., 2023). It is possible that for excretory canal outgrowth, the C1-PH-CNH module alone is sufficient for localization of MRCK-1. Technical challenges prevented observation of MRCK-1 localization in the excretory canal. Translational reporters of MRCK-1 tagged with GFP, YPet, or mNeonGreen could not be detected in the canal by confocal microscopy when expressed at endogenous levels, and overexpression of a translational reporter under a strong canal promoter caused a variety of defects that also prevented visualization of the reporter. In the future a role for the C1-PH-CNH module in MRCK-1 localization should be investigated to determine if they are required for MRCK-1 localization to the apical and/or basal membranes of the canal.

We have shown that MRCK-1 is required for outgrowth of the excretory canal when the posterior canals follow guidance cues to extend to their target positions at the tail of the worm. Other studies of MRCK function in tubulogenesis place it downstream of Cdc42-signaling for lumen formation and invasion of endothelial cells in 3D culture (Koh et al., 2008; Norden et al., 2016). In the excretory canal MRCK-1 promotes forward growth of the tube, rather than lumen formation, as *mrck-1*/MRCK-1^canal-^ mutants do form a lumen despite having extremely truncated canals. Although MRCKs in other organisms are not known to regulate biological tube outgrowth, they have been shown to function in neurite outgrowth and guidance in mammalian cell culture and the *Drosophila* visual system (Chen et al., 1999; Gontang et al., 2011; Groeger and Nobes, 2007). Many of the same molecular guidance cues that direct axon outgrowth in *C. elegans* also direct excretory canal outgrowth (Hedgecock et al., 1990; Katidou et al., 2013; Marcus-Gueret et al., 2012; McShea et al., 2013; Schmidt et al., 2009; Stringham et al., 2002) and it is well established that this also holds true for guidance factors that direct the vertebrate vascular and nervous systems (reviewed in Weinstein, 2005). Therefore, we hypothesize that MRCK-1 functions in a similar manner to promote canal and axon outgrowth and activate non-muscle myosin in response to guidance cues, although the identity of these cues and how they regulate MRCK-1 remains to be elucidated.

Mechanistically, our data suggests that MRCK-1 functions in the canal to promote phosphorylation of the regulatory light chain (RLC) MLC-4 to activate non-muscle myosin. We cannot conclude that MRCK-1 directly phosphorylates MLC-4 in the canal, but we have shown that the myosin light chain phosphatase *mel-11* does not function downstream of *mrck-1* in the canal. This demonstrates that MRCK-1 does not inhibit *mel-11* to promote MLC-4 phosphorylation, but it remains to be determined if MRCK-1 directly phosphorylates MLC-4 in this context. It is possible that MRCK-1 has additional phosphorylation targets in the canal, although the ability of the MLC-4 phosphomimetic mutants to restore canals in *mrck-1* mutants to wild type length, and our observations that MRCK-1^canal-^ and MLC-4^canal-^ mutants have the same level of canal truncation suggests this to be the primary mechanism through which MRCK-1 promotes canal outgrowth. The similar defects caused by loss of MRCK-1 and MLC-4 in the canal also suggest that MRCK-1 is the sole kinase regulating activation of non-muscle myosin in this context. MRCKs often function in parallel with another DMPK-family kinase, Rho-associated protein kinase (ROCK), to regulate non-muscle myosin (Fisher et al., 2009; Gagliardi et al., 2017; Gally et al., 2009; Groeger and Nobes, 2007; Wilkinson et al., 2005) but our findings suggest that non-muscle myosin activation in the canal is MRCK-1-driven. Previously our lab showed that knockdown of a homolog of the ROCK activator RhoA (*rho-1*) by RNAi does not cause canal truncations, suggesting that Rho/ROCK signaling is not required for excretory canal extension (Lant et al., 2015). Moreover, we propose that activation of non-muscle myosin in the canal is ROCK-independent, similar to other examples of ROCK-independent MRCK functions *in vivo* (Ando et al., 2013; Dong et al., 2002; Gomes et al., 2005; Huo et al., 2011; Marston et al., 2016).

Our observations that MLC-4/non-muscle myosin is required for canal outgrowth provides the first example of a function for non-muscle myosin in seamless tube development. Non-muscle myosin activation has been reported to function downstream of Cdc42/Rac1/Pak4 and RhoA/ROCK for lumen formation and maintenance of lumen diameter in 3D-cultured endothelial cells (Barry et al., 2016), and downstream of Rok/ROCK for salivary gland tubulogenesis in *Drosophila* (Röper, 2012). These examples of non-muscle myosin function in multicellular tubes involve the regulation of cell-cell junctions, which are absent in the seamless excretory cell. Based upon the localization of the translational MLC-4DD reporter to the growing tip of the canal, and its cell-autonomous function during canal outgrowth, it is likely that MLC-4/non-muscle myosin acts in a manner distinct from previously characterized examples in multicellular tubes. Instead, we hypothesize that MLC-4/non-muscle myosin function in the canal is similar to its function in neurites. Non-muscle myosin regulates the structure of neurites and their growth cones in different and sometimes oppositional manners, due to the expression of a variety of essential heavy chain isoforms and local activation by different kinases (Costa and Sousa, 2020). Rho/ROCK enhanced non-muscle myosin contraction of actin arcs promotes growth cone retraction (Zhang et al., 2003), while its activity can also promote retrograde flow and recycling of actin bundles in the growth cone to fuel axon extension (Medeiros et al., 2006). In *C. elegans* non-muscle myosin is required for dendrite self-avoidance (Sundararajan et al., 2019) and new growth cone formation for axon regeneration after injury (Shimizu et al., 2018). MRCK-driven activation of non-muscle myosin has been shown to promote neurite outgrowth in mammalian cell culture and axon guidance in the *Drosophila* visual system (Chen et al., 1999; Gontang et al., 2011). The function of non-muscle myosin activation downstream of MRCK-1 in the canal remains to be elucidated. We have shown that F-actin localization to the growing tip is not perturbed in *mrck-1* mutants, as F-actin is present throughout the long unlumenized cytoplasmic projections in these mutants. A similar defect is seen in neurite growth cones of the marine mollusk *Aplysia* after chemical inhibition of non-muscle myosin, where reduced actin retrograde flow and actin-bundle severing causes aberrant elongation of filopodia through extended F-actin bundles (Medeiros et al., 2006). Non-muscle myosin activity may be required in the canal to promote actin recycling in a similar manner, and the reduction of actin-bundle severing may be the underlying cause of the cytoplasmic projections seen upon loss of MRCK-1 or MLC-4.

In summary, we have defined a novel function for MRCK-1-driven activation of non-muscle myosin in seamless tube outgrowth that is independent of its canonical regulator CDC-42. Our findings show that the CRIB domain is dispensable for MRCK-1 function in the excretory canal, challenging the assumption that MRCKs require Cdc42 for proper function *in vivo*. It remains to be determined what mechanism(s) functions upstream of MRCK-1 in the canal to regulate its activation of non-muscle myosin, and how this promotes canal outgrowth. Our findings deepen our knowledge of the mechanisms of excretory canal outgrowth in *C. elegans*, which may be conserved in other seamless tubes, including vertebrate microvasculature.

## Methods

### *C. elegans* strains and maintenance

All worms were grown at 20°C on Nematode Growth Media (NGM) agar plates seeded with *Escherichia coli* OP50 bacteria (Brenner, 1974). The following strains were obtained from the *Caenorhabditis* Genetics Center (CGC) which is funded by NIH Office of Research Infrastructure Programs (P40 OK010440): FX30161, EG6699, WH556, VC141. Strain FT1722 carrying the integrated *canalp::zif-1* transgene was generously provided by Dr. Jeremy Nance (Abrams and Nance, 2021). Strain GS6603 carrying the canal-specific LifeAct::TagRFP reporter was generously provided by Dr. Daniel Shaye (Shaye and Greenwald, 2015). Strain BK205 carrying the canal-specific RAB-11::mCherry reporter was generously provided by Dr. Matthew Buechner (Mattingly and Buechner, 2011). See strain list for all strains used in this study and their genotypes.

### CRISPR/Cas9 generation of mutant alleles and transgenes

CRISPR/Cas9 genome editing was performed by direct injection of ribonucleoprotein (RNP) complex of Cas9 protein and guide RNA(s) (gRNA) as described previously (Paix et al., 2015; Popiel and Derry, 2020). All gRNAs were composed of a universal tracrRNA from IDT and target-specific crRNA(s). For deletion and substitution edits to generate *mrck-1* null, *mrck-1ΔCRIB,* and *mrck-1* KD alleles, single-stranded DNA (ssDNA) repair templates were ordered from Eurofins. For insertion of ZF1 recognition motif sequence at *mrck-1* C-terminus and *mlc-4* N-terminus, ssDNA repair templates were generated using PCR of dsDNA template followed by digestion by lambda exonuclease as described previously (Eroglu et al., 2022). All CRISPR generated alleles were confirmed by Sanger Sequencing of PCR products by The Centre for Applied Genomics (TCAG). The full list of crRNA, repair template, and primer sequences used for CRISPR/Cas9 genome editing are available in the molecular reagent table.

### Generation and/or integration of additional transgenes

The cytosolic and apical membrane canal marker *t28h11.8p::yfp::sl2::ifb-1::cfp* (*canalp::yfp::sl2::ifb-1::cfp*) was integrated as a single copy insertion from the plasmid pJA043 (a gift from Dr. Jeremy Nance) using the Mos1-mediated Single Copy Insertion (MosSCI) protocol as previously described (Frøkjær-Jensen et al., 2008).

Extrachromosomal array onEx79 (*exc-9p::gfp::mlc-4(T15D, S18D)::UTR_unc-54_*) for expression of MLC-4(DD) in the canal was generated by microinjection of plasmid pWD285 at 10 ng/µL concentration with coinjection markers pCFJ90 (Addgene Plasmid #19327) and pCFJ104 (Addgene Plasmid #19328) at 2.5 ng/µL and 5 ng/µL concentrations respectively. Microinjection was performed using a FemtoJet (Eppendorf) microinjection system with an inverted Leica DMI3000B microscope.

The plasmid pWD285 was constructed by cloning the *gfp::mlc-4::UTR_unc-54_*sequence from pML1522 (a gift from Dr. Michel Labouesse) (Gally et al., 2009) into the backbone of pBK162 (a gift from Dr. Matthew Buechner), which contains the *exc-9* promoter for expression in the canal. Q5® Site-Directed Mutagenesis (New England BioLabs, Inc.) was used to introduce nonsynonymous mutations into the *mlc-4* sequence to generate *mlc-4(T15D, S18D)*.

### Microscopy

All microscopy was performed on live worms slide mounted in M9 buffer with 5 µL tetramisole anesthetic (20 mM) on flat agarose pads (4%). All images show late L4 stage or early adult hermaphrodites unless otherwise specified. Representative confocal images of canal lengths and morphology were captured with a Leica DMI8 (TCS SP8) lighting confocal/light sheet microscope, 40x NA 1.3 water immersion (motCORR) or 63x NA 1.3 glycerol immersion objectives, 405, 448, 488, 552, and 638 nm lasers, HyD detectors, and 1x to 5x zoom. Leica LAS X software was used for image acquisition and processing. For whole worm images Leica LAS X ‘tile scanning’ with automatic stitching was used to capture the entire worm and maximum intensity projections of z-stacks were used to show the whole canal. For F-actin and GFP::MLC-4(DD) localization, the same laser power and exposure times were used within experiments, and image brightness was not altered after acquisition. For analysis of canal length and morphology fluorescent and brightfield images were viewed with a Leica DMRA2 compound microscope equipped with epifluorescence and Nomarski optics using 10x NA 0.4 or 40x NA 1.25-0.75 oil immersion objectives. Images were captured with a Hamamatsu C4742-95 digital camera using OpenLab software v5.5.2 (PerkinElmer, Inc.).

### Relative excretory canal length measurements

Relative excretory canal lengths quantify the length of one posterior branch of the canal as a ratio of the length of the worm body. Posterior canals were measured from the cell body of the excretory canal cell to the tip of the canal. For canals with cytoplasmic trailing defects, the canal was measured to the furthest lumenized part of the canal. One posterior canal for each individual was measured, chosen based upon clarity when focusing the microscope. If both canals were visible, the longest canal was measured. Length of the worm was measured from the excretory canal cell body to the tail of the worm, just past the anal pore. Measurements were made using ImageJ (NIH) software freehand trace and measurement tools. See previously published method for details (Popiel and Derry, 2020).

### Maternal embryonic/larval lethality

To quantify the maternal embryonic/larval lethality in *mrck-1* mutants and controls, L4 worms (P0) were singled out onto small NGM plates and left to lay progeny for 24 hours. After the initial 24 hours P0 individuals were moved to a fresh plate each day to facilitate counting. P0s were allowed to lay progeny (F1) for 5 days in total. After 24 hours on each plate the number of eggs laid was recorded, and the plates were monitored for 5 days each to record the number of progeny that reached adulthood (adult F1s were removed from the plate before they began to lay F2 progeny). The total number of progeny laid and the total progeny that grew to adulthood were summed across all 5 plates for each P0 to calculate the survival to adulthood of progeny.

### RNA interference by feeding

Knockdown of target genes by RNAi was performed by feeding *C. elegans* HT115 *E. coli* strain bacteria from the Ahringer library (Kamath and Ahringer, 2003). This library contains predicted gene sequences from the *C. elegans* genome cloned into the vector L4440 and transformed into HT115 *E. coli*. Bacterial colonies were grown overnight on an orbital shaker at 37°C in LB broth with Ampicillin (final [100 μg/mL]) and Tetracycline (final [10 μg/mL]). Expression of dsRNA was induced by addition of Isopropyl β-D-1-thiogalactopyranoside (IPTG) for a final concentration of 0.4 mM, and cultures remained on the orbital shaker at 37°C for an additional 4 hours after induction. Cultures were concentrated 5x before being seeded onto small NGM agar plates supplemented with carbenicillin [25 μg/mL] and IPTG [2.5 mM]. Bacterial lawns were dried and allowed to grow overnight at room temperature before worms were plated. L4 worms were placed on plates left for 48 hours to reach adulthood and lay eggs before being removed. The first generation of worms grown on RNAi were moved to lay progeny on fresh RNAi plates, and this F2 generation was assessed for phenotypes. Worms were grown for 2 generations on RNAi to allow efficient knockdown of the target gene unless the RNAi caused lethality or sterility in the first generation. Knockdown of *Y95B8A_84.g*, a non-expressed pseudogene, was used as a control for RNAi experiments (Lehner et al., 2006).

### Fluorescent intensity at canal tip

The relative fluorescent intensity of GFP::MLC-4DD at the tip of the canal was quantified using ImageJ. The canal tip and an adjacent control region were selected, and a region of interest (ROI) was created to select all pixels above threshold 20 in the image selection. Mean gray value for this selection was used as a measure of fluorescent intensity to control for differences between the area of the tip and control regions. The same tip and control regions were used to measure the fluorescent intensity of the cytoplasmic RAB-11::mCherry reporter as a control.

### Statistics

Statistical analysis was performed using GraphPad Prism 9 for all tests except Fisher’s exact test, which was performed using Rstudio (Posit Software, PBC). Statistical tests and significance levels (p-values) are specified in the figure captions. All tests performed were two-tailed. During imaging individuals were selected based upon developmental stage and orientation on the slide for ease of imaging. No data points were excluded from analysis. For canal defects in adult worms 3-4 biological replicates of n ≥ 24 individuals for each genotype were pooled to calculate the percent of the population with the phenotype. For canal defects in larval stage worms 1 biological replicate of n ≥ 19 individuals for each developmental stage was used. See sample sizes table for details of the biological replicates and sample size for each genotype.

## Acknowledgments

Many thanks to Jeremy Nance (NYU School of Medicine) and Daniel Shaye (University of Illinois Chicago) for their generous contributions of strains, plasmids, and helpful advice. Thank you to Matthew Buechner (University of Kansas) for sharing a strain and plasmid, and Michel Labouesse (Institut de Biologie Paris-Seine) for sharing a plasmid with sequence information. Thank you to Bin Yu for providing feedback and guidance for the design of CRISPR reagents. Thank you to Evan Wallace, who generated strains and preliminary data during his graduate studies that were used for this research. Thank you to Kimberly Lau of the SickKids Imaging Facility, The Hospital for Sick Children, Toronto, Canada for assistance with image acquisition on the Leica DMI8 (TCS SP8) confocal microscope. Some *C. elegans* strains were provided by the CGC, which is funded by NIH Office of Research Infrastructure Programs (P40 OD010440). This work was supported by funding from the Natural Sciences and Engineering Research Council (RGPIN-2016-06638) and the Canadian Institutes of Health Research (PJT-186205). W.B.D. is the Canada Research Chair in Animal Models of Human Disease.

