## Supplementary Information for "MRCK-1 activates non-muscle myosin for outgrowth of a unicellular tube in *Caenorhabditis elegans*"

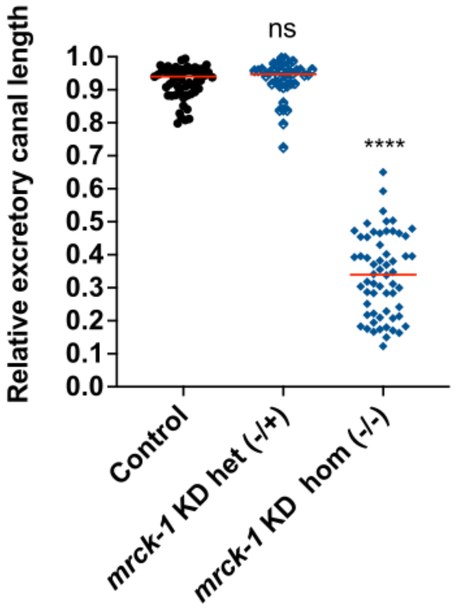


**Figure S1: *mrck-1* kinase dead allele does not function in a dominant negative manner.** There is no significant difference in the relative canal lengths of *mrck-1* KD heterozygotes (n=40) and wild type control worms (n=58). ns $p \geq 0.05$, **** $p < 0.0001$ (Mann-Whitney test versus wild type control). Red lines denote population medians.

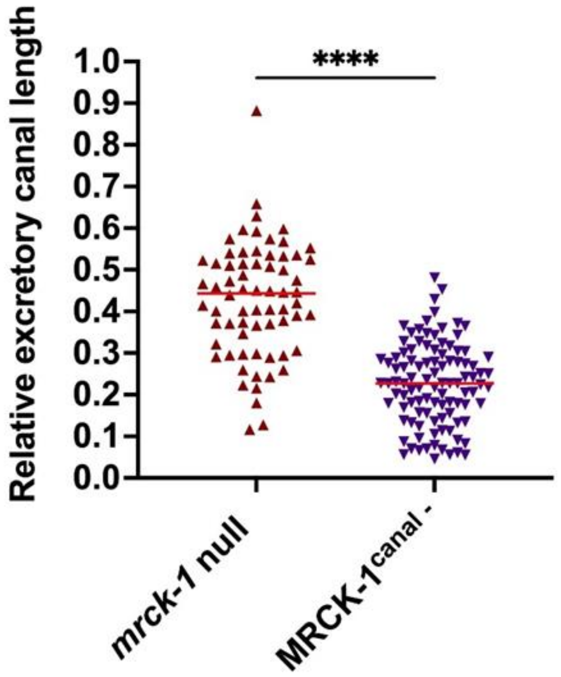


**Figure S2: Maternally contributed *mrck-1* partially suppresses canal truncations in *mrck-1* null mutants.** MRCK-1$^{canal-}$ mutants (n=102) have significantly shorter canals than *mrck-1* null mutants (n=68) due to a complete loss of maternally supplied *mrck-1*. **** $p < 0.0001$ (Mann-Whitney test). Red lines denote population medians.

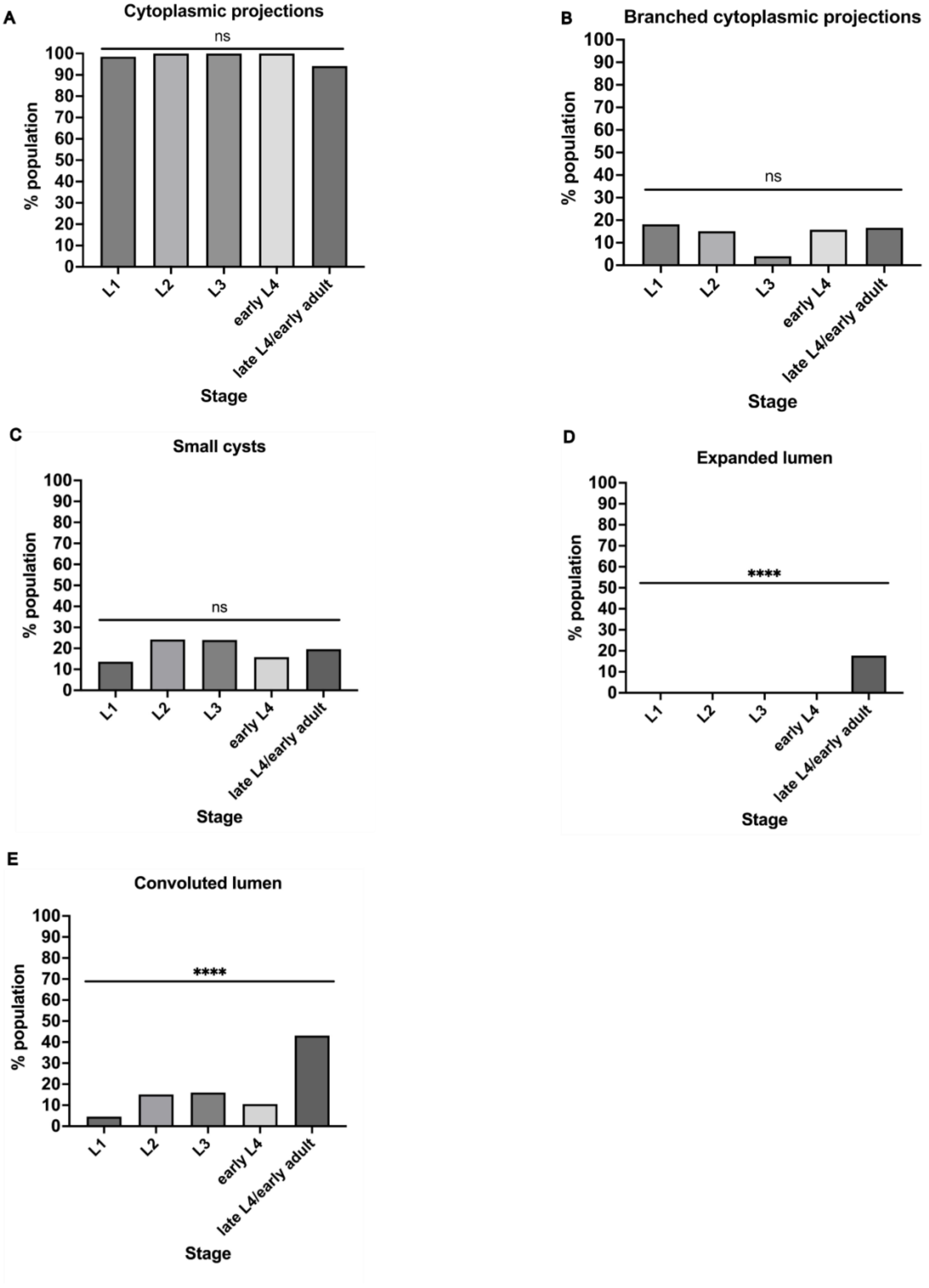
A
Cytoplasmic projections
ns
% population
Stage
L1
L2
L3
early L4
late L4/early adult
B
Branched cytoplasmic projections
ns
% population
Stage
C
Small cysts
ns
% population
Stage
D
Expanded lumen
****
% population
Stage
E
Convoluted lumen
****
% population
Stage

Figure S3: Loss of MRCK-1 in the excretory cell causes cytoplasmic projections and small cysts defects during early development. (A-C) In MRCK-1$^{canal-}$ mutants, cytoplasmic projections (single and branched) and small cysts were observed in the first larval stage and their frequency did not change significantly throughout development. (D, E) Expanded and convoluted lumens were not observed frequently in the population until maintenance during canal development (L2 stage or later), and their frequency significantly increased into adulthood. n ≥ 19 (see Materials for details). ns $p \geq 0.05$, **** $p < 0.0001$ (Fisher's exact test).

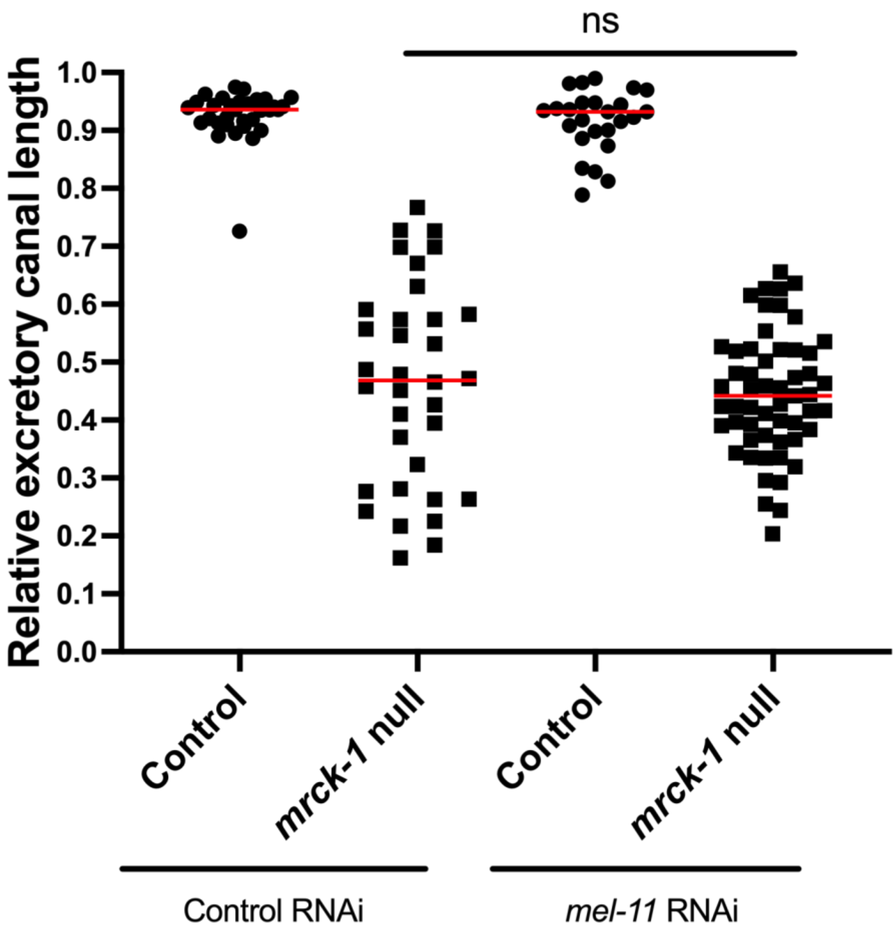

**Figure S4: Knockdown of *mel-11* does not rescue canal truncations in *mrck-1* null mutants.** There is no significant difference between canal lengths of *mrck-1* null mutants treated with *mel-11* (n=55) and control (n=34) RNAi. ns p ≥ 0.05 (Mann-Whitney test). Red lines denote population medians.

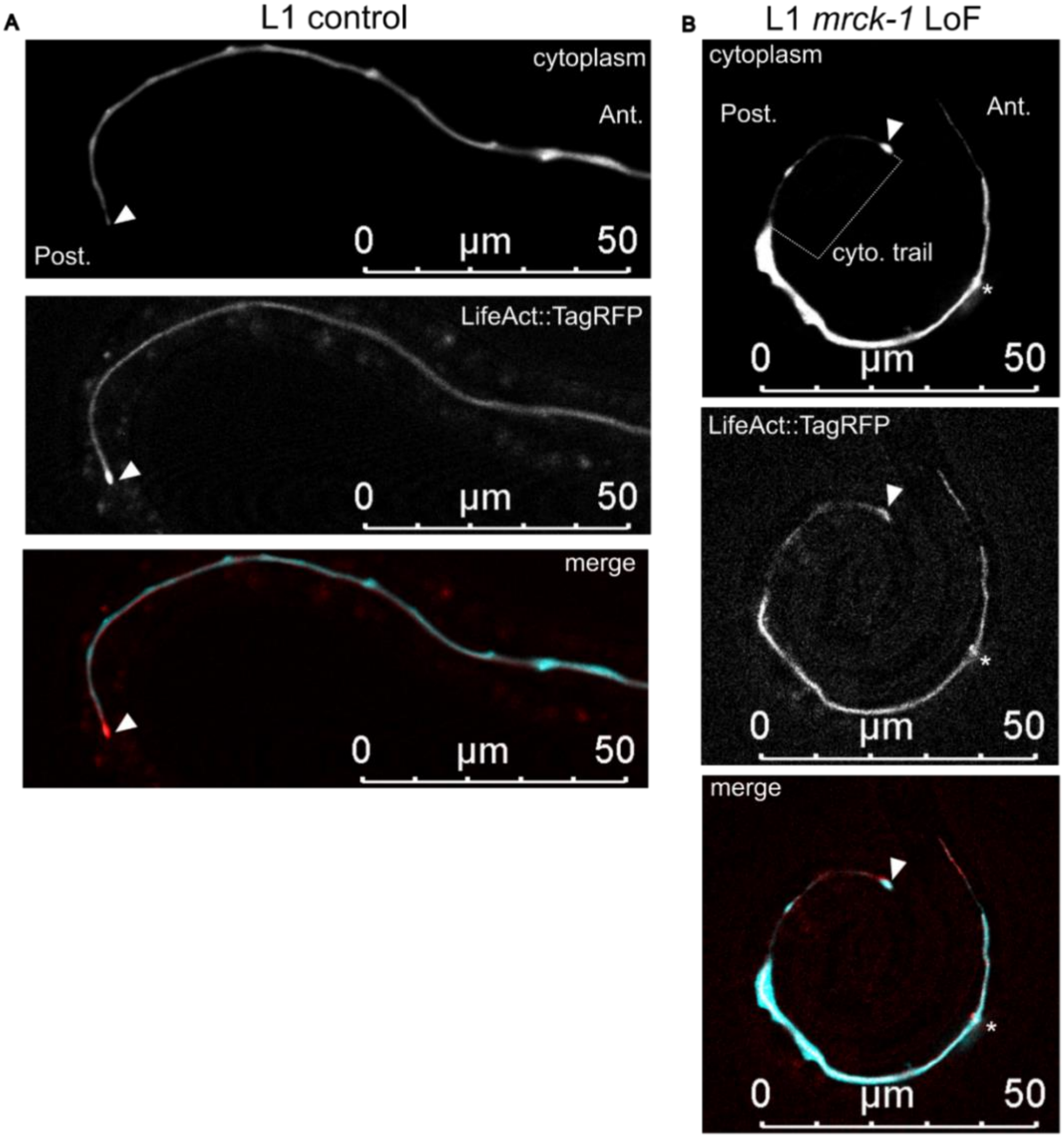


**Figure S5: Loss of *mrck-1* does not perturb F-actin localization to the growing tip of the canal during outgrowth.** In both **(A)** wild type control and **(B)** *mrck-1* LoF mutant L1 worms a marker for F-actin (LifeAct::TagRFP) is enriched at the tip of the canal (white arrowhead). Anterior (ant.) and posterior (post.) of the worms are labeled. White asterisks denote the cell body of the excretory canal.

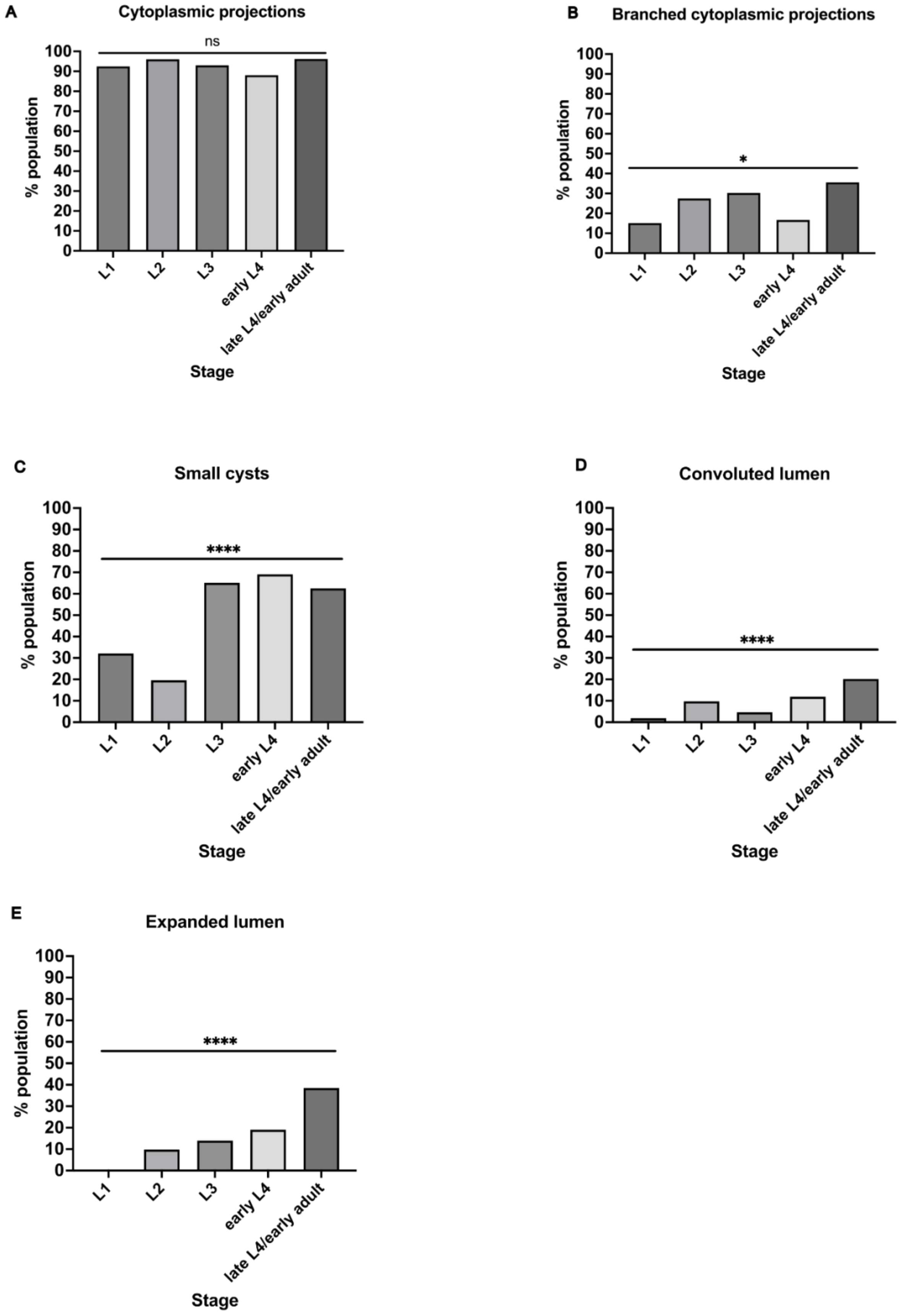
A
Cytoplasmic projections
ns
B
Branched cytoplasmic projections
*
C
Small cysts
****
D
Convoluted lumen
****
E
Expanded lumen
****
% population
Stage
L1
L2
L3
early L4
late L4/early adult

**Figure S6:** Loss of MLC-4 in the excretory cell causes cytoplasmic projections and small cyst defects during outgrowth. **(A-C)** In MLC-4$^{canal-}$ mutants, cytoplasmic projections (single and branched) and small cysts are present by L1 during canal outgrowth, while **(D, E)** convoluted and expanded lumens are observed during the maintenance phase (L2 or later). **(A)** The prevalence of cytoplasmic projections does not change during development, while **(C-E)** the prevalence of small cysts, convoluted lumens, and expanded lumens increases with age. n ≥ 36 (see Materials for details). ns p ≥ 0.05, * p < 0.05, **** p < 0.0001 (Fisher's exact test).

**Materials**

***C. elegans* strains**

| Strain name | Genotype | Source, notes |
|---|---|---|
| BK205 | *qpIs97(exc-9p::mCherry::rab-11)* | Gift from Dr. Matthew Buechner (Mattingly and Buechner, 2011). Control in Fig 1 C, D, I, K; Fig 2 B-D; Fig 3 C, D; Fig S1 |
| FX30161 | *tmC16(unc-60[tmIs1237(Pmyo-2::mCherry)])* | Ordered from the CGC (Dejima et al., 2018) |
| WD872 | *mrck-1(on167[ΔCRIB]), qpIs97/tmC16(unc-60[tmIs1237(Pmyo-2::mCherry)])* | This study |
| WD880 | *mrck-1(on171) qpIs97/tmC16* | This study. *on171* encodes *mrck-1* null allele |
| WD883 | *mrck-1(on174[K112M]) qpIs97/tmC16* | This study. *on174* encodes kinase dead allele |
| EG6699 | *ttTi5605; unc-119(ed3); oxEx1578 (eft-3p::GFP + Cbr-unc-119[+])* | Ordered from the CGC (Frøkjær-Jensen et al., 2008) |
| WD975 | *on208 (t28h11.8p::yfp::sl2::ifb-1::cfp, unc-119[+]); unc-119(ed3)* | This study |
| FT1722 | *xnIs547 (t28h11.8p::zif-1); unc-119(ed3)* | A gift from Dr. Jeremy Nance (Abrams and Nance, 2021) |

| | | |
|---|---|---|
| WD982 | *on208; unc-119(ed3); xnIs547* | This study. Control in Fig 4 A, C, E, G, I, K, M, N |
| WD990 | *on208; mrck-1(on207[mrck-1::ZF1]); xnIs54* | This study |
| WH556 | *mrck-1(ok586)/nT1(qIs51)* | Ordered from the CGC (Kumfer et al., 2010) |
| WD715 | *mrck-1(ok586)/tmC16; bgIs312 (vha-8p::gfp)* | This study |
| WD493 | *onEx79(exc-9p::gfp::mlc-4[T17D,S18D], myo-2p::mCherry, myo-3p::mCherry)* | This study |
| WD509 | *mrck-1(ok586)/nT1(qIs51); onEx79* | This study. Het was used as control in Fig 5 B |
| WD1067 | *qpIs97; onEx78(exc-9p::gfp::mlc-4[T17D,S18D], myo-2p::mCherry, myo-3p::mCherry)* | This study |
| GS6603 | *unc-119(ed3); arIs201 (glt-3p::LifeAct::TagRFP, glt-3p::CFP, unc-119[+])* | A gift from Dr. Daniel Shaye (Shaye and Greenwald, 2015). Control in Fig S5 |
| WD924 | *arIs201; mrck-1(ok586)/tmC16* | This study |
| VC141 | *zif-1(gk117)* | Ordered from the CGC (The C. elegans Deletion Mutant Consortium, 2012) |
| WD1019 | *on208; zif-1(gk117); xnIs54* | This study. Control in Fig 6 A-G; Fig 7 |
| WD1032 | *on208; zif-1(gk117), mlc-4(on228[ZF1:mlc-4]); xnIs54* | This study |

**Molecular reagents**

| Reagent name | Description | sequence (5' to 3') | Source |
|---|---|---|---|
| *mrck-1* null gRNA 1 | crRNA upstream of *mrck-1* coding sequence | GTTTGATATTTCCAGAAACA | IDT, Alt-R CRISPR-Cas9 crRNA |
| *mrck-1* null gRNA 2 | crRNA downstream of *mrck-1* coding sequence | AAATAAGTATGTAGACGAGA | IDT, Alt-R CRISPR-Cas9 crRNA |
| *mrck-1* CRIB del gRNA 1 | crRNA upstream of *mrck-1* CRIB domain sequence | TCAGGAGTGCTTCGGAGCGT | IDT, Alt-R CRISPR-Cas9 crRNA |
| *mrck-1* CRIB del gRNA 2 | crRNA downstream of *mrck-1* CRIB domain sequence | TTCGCATACTTCATCCGATA | IDT, Alt-R CRISPR-Cas9 crRNA |
| *mrck-1* kinase dead (K112M) gRNA | crRNA downstream of K112 | CGATGAAAATACTGAACAAG | IDT, Alt-R CRISPR-Cas9 crRNA |
| *mrck-1* C-term ZF1 tag gRNA | crRNA upstream of *mrck-1* STOP codon | GAGATATGTTAATATTTCAC | IDT, Alt-R CRISPR-Cas9 crRNA |
| *mlc-4* N-term ZF1 tag gRNA | crRNA upstream of *mlc-4* START codon | CAAATTTTTGCAGTAAGCCA | IDT, Alt-R CRISPR-Cas9 crRNA |
| *mrck-1* null repair template | ssDNA repair template, left homology arm 62 bp, 19 bp of sequence downstream of *mrck-1* STOP codon with edited PAM, right homology arm 60 bp | GTT CAG AAT TTG TTT TCT TTC ATT AAA TTC GTT CGT CAG GAT AGT TTG ATA TTT CCA GAA ACT TCT GCA ACT CTC TCC TTC TCG TCT ACA TAC TTA TTT TTC TCC CAA TTT CCA AAA AAT CAA AAC GAT GTA AAT CTG GTG C | Eurofins, EXTREmer |

| | | | |
|---|---|---|---|
| *mrck-1* CRIB del repair template | ssDNA repair template, left homology arm 56 bp, 14 bp mutated PAM plus sequence directly upstream of CRIB domain sequence, right homology arm 49 bp | GTA AAA ATA TTG AAT TTT TTC ATT TTA ATT ATC TAA CAA GAA ATC TCA CTG TAT GTG ACT TCG ACG CTC CGA AGC ACT CCT GAA ATT AAG CTA CCA TTT TGG TTC ATA AAA TAA TGT TG | Eurofins, EXTREmer |
| *mrck-1* kinase dead (K112M) repair template | ssDNA repair template, left homology arm 50bp, right homology arm 50bp, plus AAA to ATG substitution | CGA AAA CAC GCA GTT TCT GCT CTT TTT ACC ATT TCC CAC TTG TTC AGT ATC ATC ATC GCG TAA ATT TCA CCA ACA CCA CGC ATT CTT ACG ACG GCA ACT TCT C | Eurofins, EXTREmer |
| *mrck-1* C-term ZF1 tag repair template | dsDNA template to generate ssDNA repair template; left homology arm 150bp (with mutated PAM), last 68 bp coding sequence of *mrck-1* before stop codon, ZF recognition motif sequence (108 bp) (Armenti et al. 2014 pJN601), right homology arm 103 bp | GGGAATAAATTAATTTAAAAAATCGAG CCCGTAAATCGACACTTGCGCTACAGTA GTCATTTAAAAAACTACTGTAGTTTTCG CTAAAAGATATTTTGCCCGTAAAATAAG CTTTGCAATACGCATTCGCTGAGCTTTG TATTCACGTGAATATTAACATATCTCATT CCTAAAGCCAATTTTTTCAGGATTCACT GAACAGATCTGTAAATAACGACACAGA ATACAAAACGCGACTTTGTGATGCGTTC CGCCGTGAAGGATACTGCCCGTACAAC GACAATTGCACATATGCTCACGGACAA GATGAGCTGAGAGTTCCGAGGTAATTC TGCAACTCTCTCCTTCTCGTCTACATACT TATTTTTCTCCCAATTTCCAAAAAATCAA AACGATGTAAATCTGGTGCTATCTCCCT CTCTAGACATA | IDT, gBlocks Gene Fragments |

| | | | |
|---|---|---|---|
| *mlc-4* N-term ZF1 tag repair template | dsDNA template to generate ssDNA repair template; left homology arm 150 bp, 111 bp START codon and ZF1 recognition motif sequence, 99 bp right homolgy arm | TTT GAA CTC TTG AAT TTG AGC CTG ATC GAA CAT GGC GAA CAC ATT GGA AGT GGC TCT TTG TGG CCG TTG ACG GCG GTT TAC GGT TTT GCG GGA GGC CAT CCT CGG AAC TCT CAG CTC ATC TTG TCC GTG AGC ATA TGT GCA ATT GTC GTT GTA CGG GCA GTA TCC TTC ACG GCG GAA CGC ATC ACA AAG TCG CGT TTT GTA TTC TGT CAT TTA CTG CAA AAA TTT GAA AAT TTG CGA TTT TTG AGG AAA AAA TAA GGT CTC GAC ACG ACA AAT TTA TGT TAA ATA CTA AAA GGT GAG CGC CTT TAA AGA GTA CTG TAG TTT TAC TAA TTG GAT TTT TTT TCT CGA AAA ATC AAT GAA AAT | IDT, gBlocks Gene Fragments |
| *mrck-1::ZF1* template primer F | Forward primer for PCR to generate ssDNA repair template for *mrck-1::ZF1* transgene | GCC CGT AAA TCG ACA CTT GC | Eurofins, oligonucleotide |
| *mrck-1::ZF1* template primer R | Phosphorylated reverse primer for PCR to generate ssDNA repair template for *mrck-1::ZF1* transgene (phosphorylated strand is preferentially degraded) | [Phos]GAG AGG GAG ATA GCA CCA GA | Eurofins, oligonucleotide |
| *ZF1::mlc-4* template primer F | Forward primer for PCR to generate ssDNA repair template for *ZF1::mlc-4* transgene | GAACATGGCGAACACATTGGAAG | Eurofins, oligonucleotide |
| *ZF1::mlc-4* template primer R | Phosphorylated reverse primer for PCR to generate ssDNA repair template for ZF1::mlc-4 transgene (phosphorylated strand is preferentially degraded) | [Phos]ACA GTA CTC TTT AAA GGC GCT CAC C | Eurofins, oligonucleotide |

| | | | |
|---|---|---|---|
| pML1522 | *mlc-4p::gfp::mlc-4* | NA | A gift from Dr. Michel Labouesse (Gally et al., 2009) |
| pBK162 | *exc-9p::mCherry* | NA | A gift from Dr. Matthew Buechner |
| pWD285 | *exc-9p::gfp::mlc-4(T15D, S18D)::UTRunc-54* | NA | Generated by Evan Wallace |
| pCFJ90 | *myo-2p::mCherry::UTRunc-54* | NA | Addgene Plasmid #19327 |
| pCFJ104 | *myo-3p::mCherry::UTRunc-54* | NA | Addgene Plasmid #19328 |
| *mrck-1* null F primer | Forward primer for genotyping mrck-1 null allele | TCATCCTCCTGCATTTCCTAAC | Eurofins, oligonucleotide |
| *mrck-1* null R primer | Reverse primer for genotyping *mrck-1* null allele | GCA CAA TGA GTA CAA GAG ACA G | Eurofins, oligonucleotide |
| *mrck-1* null internal R primer | Reverse primer for genotyping *mrck-1* null allele, binds within deletion (only generates an amplicon when the wild type *mrck-1* allele is present) | GCT CGG GCC ATC CAT ATA AAT A | Eurofins, oligonucleotide |
| *mrck-1* ok586 LoF F primer | Forward primer for genotyping *mrck-1* ok586 LoF allele (external primer) | CTTTGAGGGAATCGATTGGA | Eurofins, oligonucleotide |

| | | | |
|---|---|---|---|
| *mrck-1* ok586 LoF R primer | Reverse primer for genotyping *mrck-1* ok586 LoF allele (external primer) | GAACATGACGATCCTTGGCT | Eurofins, oligonucleotide |
| *mrck-1* ok586 LoF in F primer | Forward primer for genotyping *mrck-1* ok586 LoF allele (internal primer) | GTGCTTGACATGCGTCATCC | Eurofins, oligonucleotide |
| *mrck-1* CRIB del F primer | Forward primer for genotyping *mrck-1* CRIB deletion allele | AGCAGGGAGAAAGAAGAAATTCG | Eurofins, oligonucleotide |
| *mrck-1* CRIB del R primer | Reverse primer for genotyping *mrck-1* CRIB deletion allele | CAA TGA CAC GTA AAT CGA CGC TA | Eurofins, oligonucleotide |
| *mrck-1* kinase dead (K112M) F primer | Forward primer for genotyping *mrck-1* KD (K112M) allele | TCAGCAAAGCGAAGAAGCTACG | Eurofins, oligonucleotide |
| *mrck-1* kinase dead (K112M) R primer | Reverse primer for genotyping *mrck-1* KD (K112M) allele | CTACAGCAACATTCGACGCAAC | Eurofins, oligonucleotide |
| ttTi5605 F primer | Forward primer for mosSCI insertion site ttTi5605 | TGACCCGATGAAATACGGATG | Eurofins, oligonucleotide |
| ttTi5605 R primer | Reverse primer for mosSCI insertion site ttTi5605 | GAA TTT GGC TTG TAA CGC GGA | Eurofins, oligonucleotide |
| on208 external F primer | Forward external primer for *t28h11.8p::yfp::sl2::ifb-1::cfp, unc-119(+) (on208)* transgene | TCTGGCTCTGCTTCTTCGTT | Eurofins, oligonucleotide |

| | | | |
|---|---|---|---|
| on208 internal R primer | Reverse internal primer for *t28h11.8p::yfp::sl2::ifb-1::cfp, unc-119(+) (on208)* transgene | CAATTCATCCCGGTTTCTGT | Eurofins, oligonucleotide |
| *canalp* F primer | Forward primer for *canalp::zif-1* (*xnIs547*) transgene | ATGTGGGCGTGAACAAAAA | Eurofins, oligonucleotide |
| *zif-1* R primer | Reverse primer for *canalp::zif-1* (*xnIs547*) transgene | CTA TGA ACT CGT ATT CGC T | Eurofins, oligonucleotide |
| *mrck-1::ZF1* F primer | Forward primer for *mrck-1::ZF1 (on207)* transgene | GCGTCGATTTACGTGTCATTG | Eurofins, oligonucleotide |
| *mrck-1::ZF1* R primer | Reverse primer for *mrck-1::ZF1 (on207)* transgene | GCACCAGATATGAAGGCGTTA | Eurofins, oligonucleotide |
| *zif-1(gk117)* external F primer | Forward external primer for *zif-1(gk117)* allele | CAACTTCCTTCTCTCCACTGTC | Eurofins, oligonucleotide |
| *zif-1(gk117)* external R primer | Reverse external primer for *zif-1(gk117)* allele | CAGAGAATCCACTCCGTTTGAG | Eurofins, oligonucleotide |
| *zif-1(gk117)* internal R primer | Reverse internal primer for *zif-1(gk117)* allele | CGACCACTTCCAGACAAACGAT | Eurofins, oligonucleotide |
| *ZF1::mlc-4* F external primer | Forward primer for *ZF1::mlc-4 (on228)* transgene | TTTCCTCGAGGAGTCTAGACGT | Eurofins, oligonucleotide |
| *ZF1::mlc-4* R external primer | Reverse primer for *ZF1::mlc-4 (on228)* transgene | GCTCCGGGAGCCTCGTTAATCA | Eurofins, oligonucleotide |

**Sample sizes for canal defect experiments**

| Genotype/strain | Biological replicates | n | total n (pooled) | Figure(s) data presented |
|---|---|---|---|---|

| | | | | |
|---|---|---|---|---|
| | N1 | 39 | | |
| | N2 | 31 | | |
| | N3 | 24 | | |
| Control/ BK205 | N4 | 38 | 132 | Figure 1 I, K |
| | N1 | 46 | | |
| | N2 | 31 | | |
| *mrck-1* null/WD880 | N3 | 41 | 118 | Figure 1 I, K |
| | N1 | 28 | | |
| | N2 | 47 | | |
| *mrck-1ΔCRIB*/WD872 | N3 | 51 | 126 | Figure 1 I, K |
| | N1 | 25 | | |
| | N2 | 45 | | |
| *mrck-1* KD/WD883 | N3 | 41 | 111 | Figure 1 I, K |
| | N1 | 29 | | |
| | N2 | 34 | | |
| Control/WD982 | N3 | 32 | 95 | Figure 4 C, E, G, I, K, M |
| | N1 | 32 | | |
| | N2 | 31 | | |
| MRCK-1$^{canal-}$/WD990 | N3 | 39 | 102 | Figure 4 C, E, G, I, K, M, Figure 6 B-G, Figure S3 A-E |
| L1 MRCK-1$^{canal-}$/WD990 | N1 | 66 | NA | Figure 4 N, Figure S3 A-E |
| L2 MRCK-1$^{canal-}$/WD990 | N1 | 33 | NA | Figure 4 N, Figure S3 A-E |

| | | | | |
|---|---|---|---|---|
| L3 MRCK-1$^{canal-}$/WD990 | N1 | 25 | NA | Figure 4 N, Figure S3 A-E |
| early L4 MRCK-1$^{canal-}$/WD990 | N1 | 19 | NA | Figure 4 N, Figure S3 A-E |
| | N1 | 28 | | |
| | N2 | 30 | | |
| Control/WD1019 | N3 | 33 | 91 | Figure 6 B-G |
| | N1 | 35 | | |
| | N2 | 36 | | |
| MLC-4$^{canal-}$/WD1032 | N3 | 33 | 104 | Figure 6 B-G, Figure S6 A-E |
| L1 MLC-4$^{canal-}$/WD1032 | N1 | 53 | NA | Figure 7, Figure S6 A-E |
| L2 MLC-4$^{canal-}$/WD1032 | N1 | 51 | NA | Figure 7, Figure S6 A-E |
| L3 MLC-4$^{canal-}$/WD1032 | N1 | 43 | NA | Figure 7, Figure S6 A-E |
| early L4 MLC-4$^{canal-}$/WD1032 | N1 | 43 | NA | Figure 7, Figure S6 A-E |
